# Prime editing efficiently generates W542L and S621I double mutations in two ALS genes in maize

**DOI:** 10.1101/2020.07.06.188896

**Authors:** Yuan-Yuan Jiang, Yi-Ping Chai, Min-Hui Lu, Xiu-Li Han, Qiupeng Lin, Yu Zhang, Qiang Zhang, Yun Zhou, Xue-Chen Wang, Caixia Gao, Qi-Jun Chen

## Abstract

A novel and universal CRISPR/Cas-derived precision genome-editing technology named prime editing was developed. However, low efficiency of prime editing was shown in transgenic rice lines. We reasoned that enhancing pegRNA expression would improve prime-editing efficiency. In this report, we describe two strategies for enhancing pegRNA expression. We constructed a prime-editing vector harboring two pegRNA variants for W542L and S621I double mutations in *ZmALS1* and *ZmALS2*. Compared with previous reports in rice, we achieved much higher prime-editing efficiency in maize. Our results are inspiring and provide a direction for the optimization of plant prime editors.

## Background

Despite rapid advances in genome-editing technologies, precision genome editing remains challenging [1]. Recently, a novel and universal precision genome-editing technology named prime editing was developed and tested in mammalian cells [1]. More recently, seven groups reported applications of prime editors (PE) in rice and wheat, and six groups achieved prime-edited rice lines [2-8]. Excluding the data based on enriching strategies or that are unsuitable for comparison, a total of 164 prime-edited rice lines transformed with 39 prime editors targeting 11 endogenous rice genes were generated in these previous reports [2, 3, 5, 6, 8]. The highest editing efficiencies achieved by the five groups ranged from 2.22% to 31.3% [2, 3, 5, 6, 8]. Of the total 164 prime-edited rice lines, only one line, generated by Xu et al. [6], harbors homozygous mutations; all the other lines mainly harbor chimeric mutations. These results indicate that the reported prime-editing tools require significant optimization before they are adopted for use by plant researchers. In this paper, in comparison with previous reports in rice, we achieved much higher prime-editing efficiency in maize by optimizing pegRNA expression.

## Results and discussion

We selected two maize acetolactate synthase (*ALS*) genes to test prime-editing efficiency in maize and attempted to generate maize herbicide-resistant lines harboring the P165S mutation or W542L/S621I double mutations [9] in *ZmALS1* and *ZmALS2* by prime editing (Fig. 1a). We used the maize *Ubi1* promoter to drive the expression of maize codon-optimized *PE2* and the *OsU3* and *TaU3* promoters to drive the expression of pegRNA and sgRNA, respectively. In this way, we constructed two pGreen3 binary vectors [10, 11], pZ1PE3 and pZ1PE3b, for the same P165S mutation that are based on two different strategies, PE3 and PE3b [1], respectively (Fig. 1a, b). To generate mutant maize lines harboring W542L/S621I double mutations, we used PE3b and PE3 strategies to generate the W542L and S621I mutations, respectively, and assembled binary vectors harboring two pegRNAs and two nicking sgRNA variants (Fig. 1a, b). Because of their structure of pegRNAs, a hairpin can potentially form between the primer-binding site (PBS) and the protospacer of a pegRNA (Fig. 1a), possibly weakening pegRNA activity. Considering this possibility, we reasoned that enhancing pegRNA expression would improve the editing efficiency. We used two strategies to enhance pegRNA expression: doubling the number of pegRNA expression cassettes and using two promoter systems to drive pegRNA expression, together with tRNA, ribozyme, and Csy4 RNA processing systems [12, 13] as appropriate (Fig. 1b). In this way, we generated two prime editors named pZ1WS-Csy4 and pZ1WS (Fig. 1b). We also generated two additional prime editors, p4xZ1PE3-Csy4 and p4xZ1PE3b-Csy4, for comparison with pZ1PE3 and pZ1PE3b, respectively. The p4xZ1PE3-Csy4 and p4xZ1PE3b-Csy4 editors have structures similar to that of pZ1WS-Csy4 but harbor redoubled the pegRNA and sgRNA expression cassettes for the P165S mutation. The pZ1WS vector was constructed to use as a control of pZ1WS-Csy4 for comparing the non-Csy4 system with the Csy4 system. We transformed maize with these six PE vectors via the Agrobacterium-mediated method. No transgenic lines were obtained with the three Csy4 vectors. Almost simultaneously, we noticed that transformations with 13 additional pGreen3 or pCambia binary vectors harboring the Csy4-P2A-ZsGreen fusion gene, including 1 maize and 12 Arabidopsis transformations, were unsuccessful, suggesting that Csy4 protein severely affects Agrobacterium-mediated transformation in, at least, these two plant species.

**Fig. 1.**
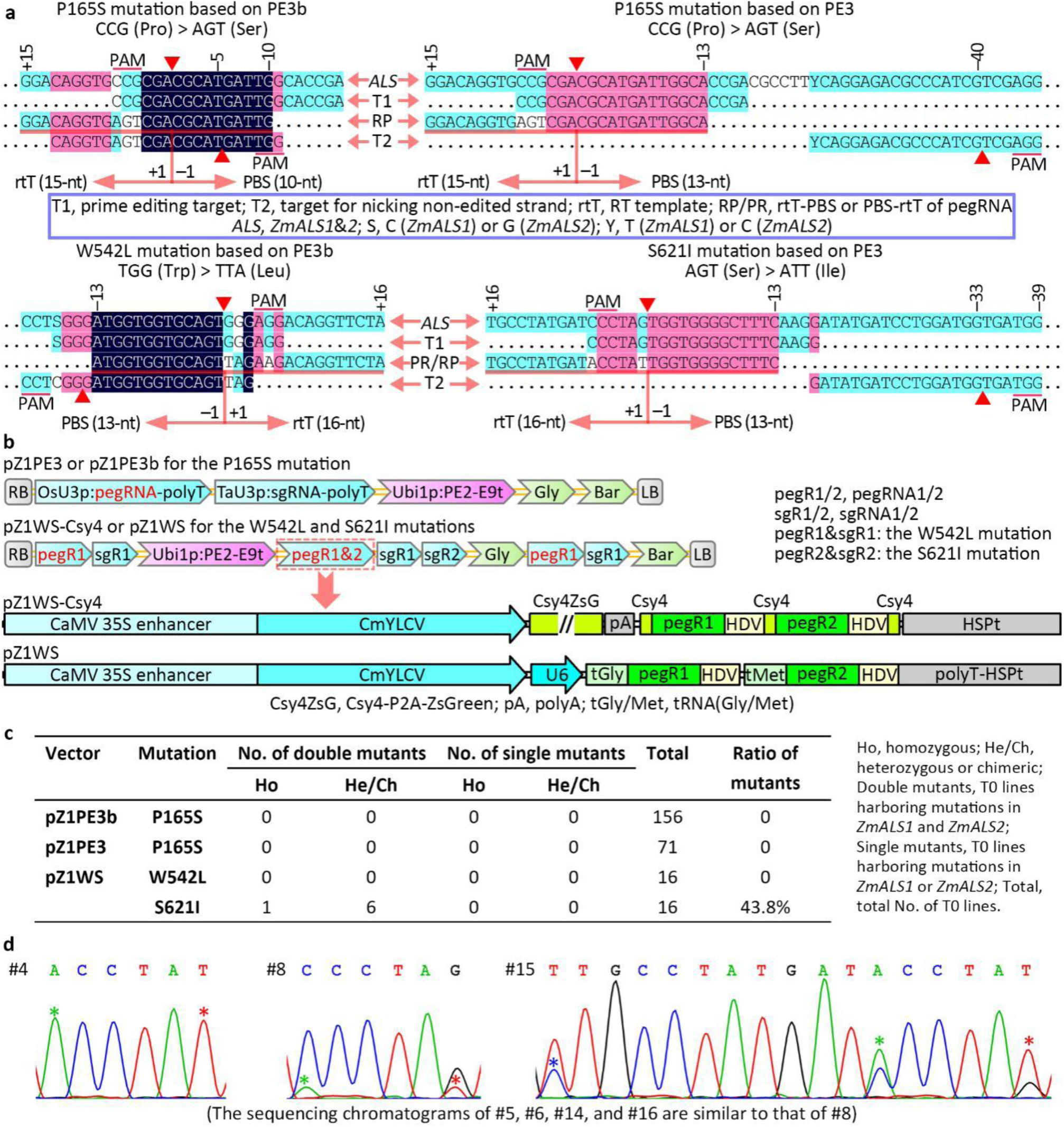
Efficiency of prime editing in maize. **a** Sequences and their relationships of the two *ALS* genes, the pegRNA targets, the pegRNA RT templates, the pegRNA primer-binding sites (PBS), and the sgRNA targets for nicking the non-edited strands used for the generation of 4 maize prime-editing vectors. The PAMs, nicking sites, RT template and PBS lengths and introduced mutations are indicated. **b** T-DNA structures of the 4 PE vectors. The pZ1WS and pZ1WS-Csy4 vectors were designed for editing two targets. Each double-PE vector included two expression cassettes for each of the two pegRNAs or sgRNAs. Two strategies were used to express peg1&2: one was based on the Csy4 RNA processing system, and the other was based on tRNA and HDV ribozyme RNA-processing systems integrated with two drivers, polymerase II (35S enhancer-CmYLCV) and III (shortened U6-26) promoters. The structures of the pegRNA and sgRNA expression cassettes in pegR1-sgR1 and pegR2-sgR2 are the same as those in pZ1PE3/3b. The expression of two sgRNAs in sgR1-sgR2 is driven by OsU3p and TaU3p. **c** Prime-editing efficiencies based on direct sequencing of the PCR products. The transgenic lines transformed with pZ1WS-Csy4 were not obtained. **d** Sequencing chromatograms of the PCR fragments from prime-edited plants transformed with pZ1WS. Double peaks represent heterozygous or chimeric mutations, and an asterisk indicates a mutation induced by PE. The first mutation in #15 is the unwanted mutation introduced by the pegRNA scaffold.

To confirm the targeted mutations, we amplified fragments spanning the target sequences from the genomic DNA of the transgenic lines and detected the mutations by direct sequencing of the PCR fragments. We observed no editing in the 71 and 156 lines transformed with pZ1PE3 and pZ1PE3b, respectively. Fortunately, we observed that 7 of 16 lines transformed with pZ1WS harbored the S621I edit (Fig. 1c, d). Interestingly, one line displayed homozygous mutations in both *ZmALS1* and *ZmALS2*.

We cloned the PCR products from the 6 lines harboring chimeric or heterozygous mutations and sequenced ∼86 clones for each PCR fragment (Additional file 1: Table S1). Interestingly, we observed the W542L edits in some cloned fragments, suggesting that low-frequency mutations could not be displayed in sequencing chromatograms (Additional file 1: Table S1). To further analyze the low-frequency W542L and/or S621I mutations, we cloned the PCR fragments from the line (#4) harboring homozygous mutations and an additional line (#3) harboring no mutations based on direct sequencing of the PCR fragments. We observed the S621I edits in some cloned fragments from line #3, further demonstrating that low-frequency mutations could not be reflected by direct sequencing of the PCR products (Additional file 1: Table S1). Three cloned fragments harbored W542L and S621I double mutations, suggesting that PE can simultaneously edit two nonallelic targets in a cell. We also cloned 6 PCR fragments from three pZ1PE3 and three pZ1PE3b transgenic lines and sequenced the cloned PCR fragments but found no mutations in a total of 213 *ZmALS1* and 196 *ZmALS2* clones (Additional file 1: Table S1).

We observed two types of undesired byproducts from the sequencing results of the cloned PCR fragments. One type was derived from the pegRNA scaffold, which acts as extended RT templates and thus can introduce unwanted edits; the other was involved only in editing of multiple nucleotides and was derived from unbiased and well-balanced double-strand repair (not single-strand-biased repair) of multiple mismatches in heteroduplex DNA, which led to incomplete editing of multiple nucleotides (Fig. 2a). In addition, the two mechanisms for forming these two types of byproducts sometimes overlapped (Fig. 2a). We observed high frequencies of byproducts at the S621 target: 23.8% and 7.2% were derived from the pegRNA scaffold and double-strand even DNA repair, respectively (Fig. 2b). For the W542 target, the frequencies of both types of byproducts remain to be investigated because of the limited number of cloned PCR fragments harboring W542L edits. Since T0 line #4 harbors homozygous mutations in the two *ALS* genes and because T0 line #15 harbors obvious pegRNA scaffold-derived byproducts (Fig. 1d), we excluded these two lines to perform an analysis of the two types of byproducts that was as unbiased as possible. The recalculated frequencies of the byproducts for the S621I edits were 18.1% and 15.3% for the pegRNA scaffold-derived and double-strand even DNA repair-derived byproducts, respectively (Additional file 1: Table S1), indicating that frequencies of byproducts excluding the two lines were still high. Surprisingly, we observed no indel-type byproducts, suggesting that indels are infrequently induced by PE in maize.

**Fig. 2.**
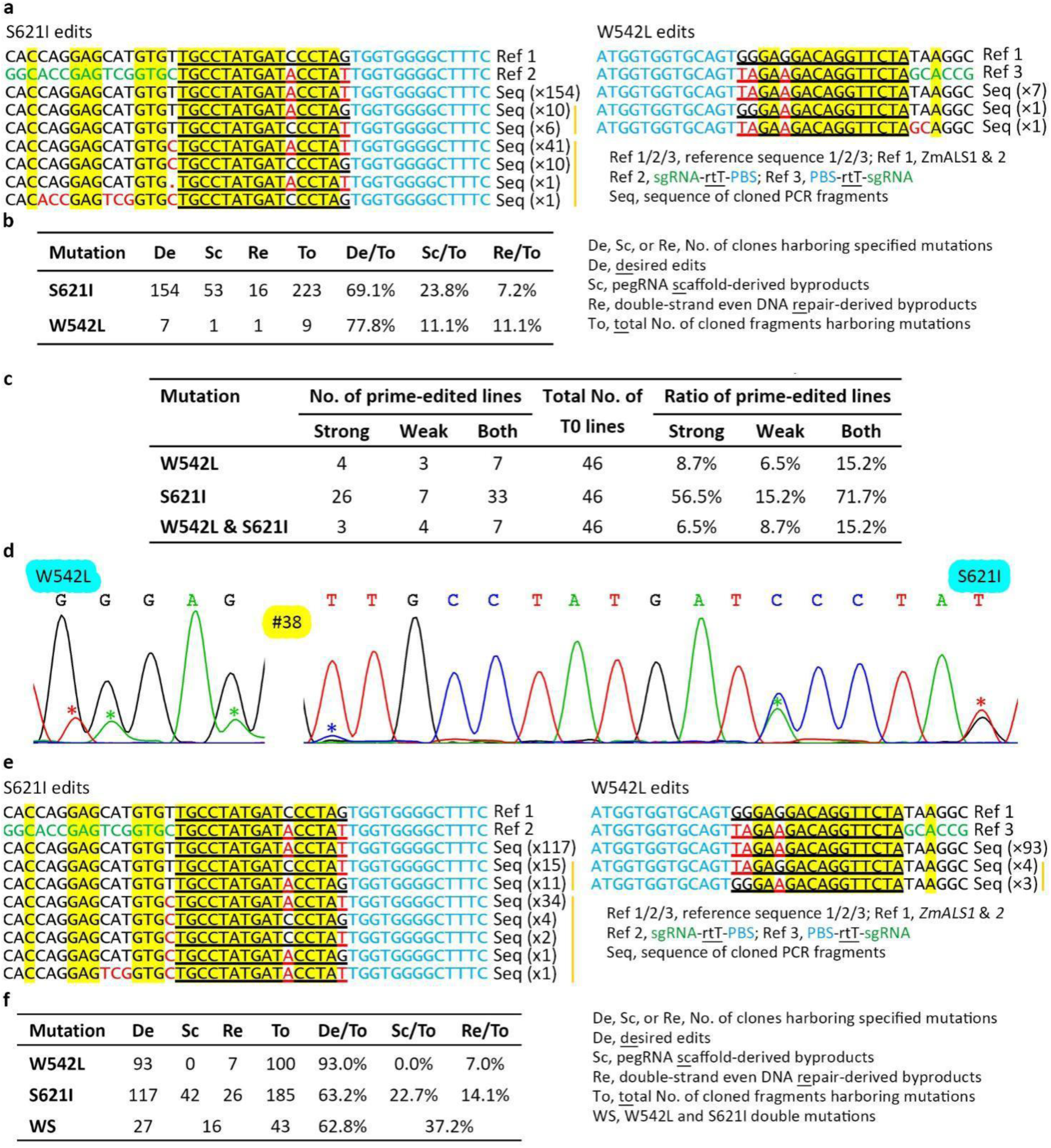
Desired edits and unwanted byproducts in the two transformations with pZ1WS. **a** Desired edits and unwanted byproducts at the two target sites in the first transformation. Only partial sgRNA sequences are shown. The homologous sgRNA-rtT sequences shared by all the aligned sequences are shaded in yellow, and the mutated nucleotides are indicated by red letters. The number of cloned PCR fragments harboring the same edits is indicated in parentheses, and a vertical line indicates the same type of byproduct. For convenience, the byproducts derived from the two mechanisms were assorted into pegRNA scaffold-derived byproduct category. **b** Summary of the desired edits and byproducts from the first transformation according to sequences of the cloned PCR fragments. **c** Prime-editing efficiency achieved in the second transformation. Strong and Weak, T0 lines harboring strong and weak peak signals for the edits, respectively, in the sequencing chromatograms. Both, total No. of lines harboring strong and weak edits. **d** Sequencing chromatograms from a prime-edited line harboring W542L edits and obtained in the second transformation. Double peaks represent heterozygous or chimeric mutations, and an asterisk indicates a mutation induced by PE. The first mutation in the S621 target is the pegRNA scaffold-derived byproducts. **e** Desired edits and unwanted byproducts at the two target sites in the second transformation. **f** Summary of the desired edits and byproducts in the second transformation.

To further determine the low-frequency mutations and byproducts, we analyzed all 243 lines transformed with the three PE vectors by next-generation sequencing (NGS) of PCR amplicons. Of the 227 T0 lines transformed with pZ1PE3 or pZ1PE3b, we found only one line transformed with pZ1PE3 harboring the low frequency (0.07%, 21/30306) P165S edits. In contrast, we found that 12 of the 16 T0 lines transformed with pZ1WS harbored the S621I edits, and 6 of the 16 lines also harbored the W542L edits (Additional file 1: Table S2). For the S621 target, the average frequency of pegRNA scaffold-derived and double-strand even DNA repair-derived byproducts based on NGS was 17.5% and 8.5%, respectively; when lines #4 and #15 were excluded, the average was 12.1% and 16.2%, respectively (Additional file 1: Table S2). We observed no byproducts for the W542L edits from NGS, possibly owing to the low frequency of the desired edits, which affected the analysis of the low-frequency byproducts.

We repeatedly transformed maize with pZ1WS after we observed that our first transformation did not generate many transgenic lines. In the second transformation, we obtained 46 transgenic lines. We detected targeted mutations induced by prime editing, and the results demonstrated again that the prime-editing efficiency of pZ1WS was much higher than that in previous reports (Fig. 2c and Additional file 1: Fig. S1). Interestingly, we observed the signals of the W542L edits in the sequencing chromatograms (Fig. 2d and Additional file 1: Fig. S1). These results suggest that we can obtain homozygous mutants harboring W542L and S621I double mutations in both *ZmALS1* and *ZmALS2* in the progeny of some lines, such as line #38 (Additional file 1: Fig. S1). In addition, we achieved two additional lines harboring S621I homozygous mutations in the two *ALS* genes (Additional file 1: Fig. S2), and these two lines, together with the line described above, will help to generate homozygous double mutants harboring W542L and S621I double mutations in the progeny. Not surprisingly, we again observed pegRNA scaffold-derived byproducts in the sequencing chromatograms (Additional file 1: Fig. S1). We cloned PCR products from the 4 lines harboring strong W542L edit signals in the chromatograms (Additional file 1: Fig. S1) and sequenced ∼106 clones for each PCR fragment (Fig. 2e, f and Additional file 1: Table S3). The results indicated that 11.5% of the cloned fragments from line #41 harbored S621I edits, although direct sequencing of PCR fragments from this line displayed weak S621I edit signals in the chromatograms (Additional file 1: Fig. S1 and Table S3). We detected no pegRNA scaffold-derived byproducts due to the W542L edits (Fig. 2e, f and Additional file 1: Table S3), suggesting that a low frequency of pegRNA scaffold-derived byproducts was produced for this target and that the frequency of the byproducts depended on the targets. Of 242 cloned PCR fragments harboring edits, 17.8% (43/242) harbored W542L and S621I double mutations in *ZmALS1* or *ZmALS2* (Fig. 2f), demonstrating again that PE can simultaneously edit two nonallelic targets in a cell. For the W542L edits, the protospacer of the sgRNA nicking non-edited strand fully matched the edited *ZmALS1* target but a mismatch was adjacent to the PAM of the edited *ZmALS2* target (Fig. 1a); therefore, PE2 was used to edit *ZmALS2*, and PE3b was used to edit *ZmALS1*. The editing frequency of the W542 target in *ZmALS1* and *ZmALS2* was 7.1% and 39.7%, respectively (Additional file 1: Table S3), suggesting that PE2 has a much higher editing efficiency than PE3b.

Since the prime-editing efficiency at the W542 target was much lower than that at S621 although the same two strategies were used to enhance the expression of the two pegRNA genes, the results suggested that improving prime-editing efficiency requires the cooperation of optimal pegRNA design with enhanced pegRNA expression. To determine the individual contributions of the two strategies for enhancing pegRNA expression, we generated seven rice prime editors and tested their editing efficiencies in rice protoplasts (Additional file 1: Fig. S3a). The results indicated that the strategy based on CaMV35S-CmYLCV-U6 composite promoter significantly improved the prime-editing efficiencies for three of four pegRNAs (Additional file 1: Fig. S3b and Table S4). However, the strategy based on redoubling the number of pegRNA cassettes had no contributions to improving prime-editing efficiency (Additional file 1: Fig. S3c and Table S4). It remains to be investigated whether doubling or redoubling the number of pegRNA expression cassettes contributes to improving prime-editing efficiency in stably transformed rice plants.

Collectively, we obtained sufficiently high efficiency for the prime editing of maize genes, possibly owing to the optimized pegRNA expression. In addition, we observed a high frequency of pegRNA scaffold-derived byproducts, and to reduce the number of byproducts, we proposed a rule of termination for the design of pegRNAs: from one to three nucleotides (C, GC, or TGC) of the pegRNA scaffold adjoining the RT template can be used as termination signals in genomic DNA for RT templates.

## Conclusions

Compared with previous reports in rice, our study achieved much higher prime-editing efficiency in maize: 53.2% (33/62) and 6.5% (4/62) transgenic lines harboring S621I and W542L mutations, respectively, in *ZmALS1* and/or *ZmALS2*; 4.8% (3/62) and 4.8% (3/62) lines harboring homozygous S621I mutations and S621I/W542L double mutations, respectively, in the two *ALS* genes. We also observed a high frequency of pegRNA scaffold-derived byproducts, and to potentially overcome this defect, we proposed a rule for designing pegRNAs. Our results are inspiring and provide a new reference standard for future optimization of prime editors. In addition, the pZ1WS prime editor generated in this study is useful for breeding transgene-free maize lines harboring W542L and S621I double mutations in the two ALS genes, which confer resistance to multiple herbicides functioning as ALS inhibitors.

## Methods

All primers used in this study are listed in Additional file 1: Table S5, and the sequences of the PE2 and pegRNA expression cassettes are listed in Additional file 1: Supplemental material. Vectors described in this study together with their annotated sequences are available from Addgene and/or MolecularCloud (GenScript).

### Vector construction

We replaced the *Xba*I-*Sbf*I fragment of pUC57-dCas9 [14] with a synthetic fragment, resulting in the generation of pUC57-PE2P1. We replaced the *Mlu*I and *Sac*I fragment of pUC57-PE2P1 with another synthetic fragment, resulting in the generation of pUC57-PE2. We modified the maize Ubi1 promoter in pG3R23-U3Ub [11] to disrupt the first of the two *Pst*I sites, the first of the four *Xba*I sites (the last 3 *Xba*I sites are blocked by Dam methylation), the *Nco*I site, and the *Eco*RI site in the promoter. We replaced the *Xba*I and *Sac*I fragment of modified pG3R23-U3Ub with the *Xba*I and *Sac*I fragment of pUC57-PE2, resulting in the generation of pG3R23-PE2U3A. We replaced the *Eco*RI-*Sac*II fragment of pG3R23-PE2U3A with the *Eco*RI-*Sac*II fragment of pG3GB411 [11], resulting in the generation of pG3GB-PE2U3A. To generate the final PE binary vectors harboring a single pegRNA, we used the Golden Gate method to insert a synthetic pegRNA-OsU3t-TaU3p-sgRNA fragment into the two *Bsa*I sites of pG3GB-PE2U3A. In this way, we generated pZ1PE3 and pZ1PE3b.

We replaced the *Asc*I-*Pac*I fragment of pL21-iCre [11] with a synthetic fragment of 35S-CmYLCV-Csy4-P2A-ZsGreen-polyA-Csy4-BB-HDV-HSPt, resulting in the generation of pL2L1-35Csy4. We replaced the *Xba*I-*Eco*RI fragment of pL2L1-35Csy4 with a synthetic fragment of U6-tGly-BB-HDV-HSPt, resulting in the generation of pL2L1-35C-U6. We replaced the *Hin*dIII-*Eco*RI fragment of pR14-BWM [11] with an insert prepared by annealing two oligos, oHEASmE-F/R, resulting in the generation of pR1R4-HEAS. We replaced the *Eco*RI-*Aar*I fragment of pR1R4-HEAS with the *Eco*RI-*Spe*I fragment of synthetic OsAct1p-CP4EPSPS-NOSt, resulting in the generation of pR1R4-Gly. We amplified the U3p-BB-sgRNA cassette from pG3R23-PE2U3A with primers OsU3p-AsF and TaU3t-EcR, purified the PCR products, digested them with *Asc*I and *Eco*RI, and allowed them to ligate with *Asc*I and *Eco*RI-digested pR1R4-Gly and pL43-U3-Bar, which resulted in the generation of pR1R4-U3G.0 and pL4L3-U3B.0, respectively.

We generated two binary vectors harboring four pegRNAs and four sgRNA cassettes in two steps. First, we used Golden Gate method to clone synthetic fragments harboring pegRNAs and/or sgRNAs into the destination vector pG3R23-PE2U3A and the three entry vectors including pL2L1-35Csy4 or pL2L1-35C-U6, pR1R4-U3G.0, and pL4L3-U3B.0 (Additional file 1: Supplemental material). Second, we used MultiSite Gateway technology to assemble the final PE vectors. In this way, we generated pZ1WS-Csy4 and pZ1WS. We constructed p4xZ1PE3-Csy4 and p4xZ1PE3b-Csy4 harboring redoubled pegRNA and sgRNA expression cassettes to generate the P165S mutation in a manner similar to the generation of pZ1WS-Csy4.

To generate rice prime editors harboring a single or two pegRNAs, we replaced the *Hin*dIII-*Spe*I fragment of pG3R23-PE2U3A with an insert prepared by annealing two oligos, oiSce-HSF/-HSR, resulting in the generation of pG3R23-PE2-dA. We replaced the *Bas*I-*Bsa*I fragment of pG3R23-PE2-dA with the *Hin*dIII-*Eco*RI fragment of pL2L1-35C-U6, resulting in the generation of pG3R23-PE2-35C. We replaced the *Hin*dIII-*Spe*I fragment of pG3R23-PE2-dA with a *Hin*dIII-*Spe*I fragment of OsU3p-tGly-BB-HDV-TaU3t, resulting in the generation of pG3R23-PE2-U3A.2. We replaced the *Eco*RI-*Sac*II fragments of pG3R23-PE2-35C and pG3R23-PE2-U3A.2 with the *Eco*RI-*Sac*II fragment of pHSE401, resulting in the generation of pG3H-PE2-35C and pG3H-PE2-U3. To generate the six rice PE vectors harboring a single or two pegRNAs, we used the Golden Gate method to insert synthetic fragments (Additional file 1: Supplemental material) into the two *Bsa*I sites of pG3H-PE2-35C or pG3H-PE2-U3. In this way, we generated pU3-ALS-S1, p35C-ALS-S1, pU3-GAPDH, p35C-GAPDH, p2xU3-ALS-WS, and p35C-ALS-WS.

To generate the rice prime editor harboring eight pegRNAs, we replaced the *Asc*I-*Eco*RI fragments of pR1R4-U3G.0 and pL4L3-U3B.0 with an *Asc*I-*Eco*RI fragment of OsU3p-tGly-BB-HDV-TaU3t, resulting in the generation of pR1R4-U3G.2 and pL4L3-U3B.2. We replaced the *Eco*RI-*Sac*II fragment of pL4L3-U3B.2 with the *Eco*RI-*Sac*II fragment of pHSE401, resulting in the generation of pL4L3-U3H.2. We generated the rice PE binary vector harboring eight pegRNAs in two steps. First, we used the Golden Gate method to clone synthetic fragments harboring two pegRNAs (Additional file 1: Supplemental material) into the destination vector pG3R23-PE2U3A.2 and the three entry vectors including pL2L1-35C-U6, pR1R4-U3G.2, and pL4L3-U3H.2. Second, we used MultiSite Gateway technology to assemble the final PE vector pALS-WSx4.

### Maize transformation and analysis of prime editing

We separately transformed the six PE vectors, pZ1PE3, pZ1PE3b, p4xZ1PE3-Csy4, p4xZ1PE3b-Csy4, pZ1WS-Csy4, and pZ1WS, into the engineered Agrobacterium strain LBA4404/pVS1-VIR2 to generate strains harboring the ternary vector system [11]. We used these strains separately to transform maize ND73, a B73-derived inbred line, according to published protocols [11].

To analyze the P165S mutations in *ZmALS1* and *ZmALS2*, we simultaneously amplified two fragments spanning the target site in the two *ALS* genes from genomic DNA of the transgenic lines using PCR with ALS1&2P-F/R primers. We then submitted the purified PCR products to direct sequencing with the ALS1&2P-F primer. To analyze the W542L and S621I mutations in *ZmALS1* and *ZmALS2*, we simultaneously amplified two fragments spanning the two target sites in the two *ALS* genes using PCR with the ALS1&2WS-F/R primers and submitted the purified PCR products to direct sequencing with the ALS1&2WS-F2 primer. Since we designed the primers according to homologous sequences of *ZmALS1* and *ZmALS2*, the PCR fragments contained both *ZmALS1* and *ZmALS2* sequences. Since *ZmALS1* and *ZmALS2* are highly homologous, the coexistence of two PCR fragments can be determined according to the double peaks in the sequencing chromatograms. Similarly, chimeric and heterozygous mutations can also be determined according to double peaks. For sequencing cloned PCR fragments, we picked ∼86 or ∼106 colonies from each transformation and submitted them for Sanger sequencing. We also used a Hi-TOM assay [15] with a 0.5% threshold to analyze the mutations in T0 transgenic plants. To analyze the P165S, W542L, and S621I mutations in *ZmALS1* and *ZmALS2*, we simultaneously amplified two fragments spanning the target site in the two *ALS* genes from genomic DNA of the transgenic lines using PCR with primers ALS1&2P-NGSF/R, ALS1&2W-NGSF/R, and ALS1&2S-NGSF/R, respectively.

### Rice protoplast transfection and analysis of prime editing

The plasmids were transferred into protoplasts by PEG-mediated methods and the transfected protoplasts were incubated at 26 °C for 48 hours. To analyze of prime editing, the genomic DNA was extracted and the primers listed in the additional file (Additional file 1: Table S5) were used to amplify target fragments for deep sequencing. For each target site, sequencing fragment was repeated three times using genomic DNA extracted from three independent protoplast samples.

## Supplementary information

**Additional file 1: Figure S1**. Sequencing chromatograms from 7 prime-edited lines harboring W542L edits. **Figure S2**. Sequencing chromatograms from 2 prime-edited lines harboring homozygous S621I edits. **Figure S3**. Prime-editing efficiency in rice protoplasts for pegRNAs based on different expression strategies. **Table S1**. Edits and byproducts revealed from cloned PCR fragments. **Table S2**. Analysis of mutations in T0 transgenic plants by NGS with a 0.5% threshold. **Table S3**. Edits and byproducts from the 4 additional lines. **Table S4**. Prime-editing efficiency in rice protoplasts analyzed by NGS. **Table S5**. Sequences of primers, targets, and rtT-PBS of the pegRNAs.

## Supplemental material

Sequences of the PE2 and pegRNA expression cassettes.

## Acknowledgments

The transgenic maize lines were created by the Maize Functional Genomic Platform of China Agricultural University. We thank our colleagues from the platform for help with the maize transformations.

## Authors’ contributions

QJC, YZhou, CG, and XCW conceived and designed the research. YYJ, YPC, MHL, XLH, QL, YZ, and QZ conducted the experiments and analyzed the data. QJC, CG, YZhou, and XCW wrote the paper. All authors read and approved the final version.

## Availability of data and materials

The vectors described in this study together with their annotated sequences are available from Addgene and/or MolecularCloud (GenScript).

## Ethics approval and consent to participate

Not applicable.

## Consent for publication

Not applicable.

## Author details

Not applicable.

## References

1. Anzalone AV, Randolph PB, Davis JR, Sousa AA, Koblan LW, Levy JM, Chen PJ, Wilson C, Newby GA, Raguram A, Liu DR: Search-and-replace genome editing without double-strand breaks or donor DNA. Nature 2019, 576:149–157.

2. Xu R, Li J, Liu X, Shan T, Qin R, Wei P: Development of a plant prime editing system for precise editing in the rice genome. Plant Communications 2020:100043.

3. Hua K, Jiang Y, Tao X, Zhu JK: Precision genome engineering in rice using prime editing system. Plant Biotechnol J 2020.

4. Butt H, Rao GS, Sedeek K, Aman R, Kamel R, Mahfouz M: Engineering herbicide resistance via prime editing in rice. Plant Biotechnol J 2020.

5. Lin Q, Zong Y, Xue C, Wang S, Jin S, Zhu Z, Wang Y, Anzalone AV, Raguram A, Doman JL, et al: Prime genome editing in rice and wheat. Nature Biotechnology 2020.

6. Xu W, Zhang C, Yang Y, Zhao S, Kang G, He X, Song J, Yang J: Versatile Nucleotides Substitution in Plant Using an Improved Prime Editing System. Molecular Plant 2020.

7. Tang X, Sretenovic S, Ren Q, Jia X, Li M, Fan T, Yin D, Xiang S, Guo Y, Liu L, et al: Plant Prime Editors Enable Precise Gene Editing in Rice Cells. Molecular Plant 2020.

8. Li H, Li J, Chen J, Yan L, Xia L: Precise Modifications of Both Exogenous and Endogenous Genes in Rice by Prime Editing. Molecular Plant 2020.

9. Kawai K, Kaku K, Izawa N, Shimizu T, Fukuda A, Tanaka Y: A novel mutant acetolactate synthase gene from rice cells, which confers resistance to ALS-inhibiting herbicides. Journal of Pesticide Science 2007, 32:89–98.

10. Zhang Y, Zhang Q, Chen QJ: Agrobacterium-mediated delivery of CRISPR/Cas reagents for genome editing in plants enters an era of ternary vector systems. Sci China Life Sci 2020.

11. Zhang Q, Zhang Y, Lu MH, Chai YP, Jiang YY, Zhou Y, Wang XC, Chen QJ: A Novel Ternary Vector System United with Morphogenic Genes Enhances CRISPR/Cas Delivery in Maize. Plant Physiol 2019, 181:1441–1448.

12. Zhang Q, Xing HL, Wang ZP, Zhang HY, Yang F, Wang XC, Chen QJ: Potential high-frequency off-target mutagenesis induced by CRISPR/Cas9 in Arabidopsis and its prevention. Plant Mol Biol 2018, 96:445–456.

13. Cermak T, Curtin SJ, Gil-Humanes J, Cegan R, Kono TJY, Konecna E, Belanto JJ, Starker CG, Mathre JW, Greenstein RL, Voytas DF: A Multipurpose toolkit to enable advanced genome engineering in plants. Plant Cell 2017, 29:1196–1217.

14. Xing HL, Dong L, Wang ZP, Zhang HY, Han CY, Liu B, Wang XC, Chen QJ: A CRISPR/Cas9 toolkit for multiplex genome editing in plants. BMC Plant Biol 2014, 14:327.

15. Liu Q, Wang C, Jiao X, Zhang H, Song L, Li Y, Gao C, Wang K: Hi-TOM: a platform for high-throughput tracking of mutations induced by CRISPR/Cas systems. Sci China Life Sci 2019, 62:1–7.

