## Supplemental Figures and Tables for "Prime editing efficiently generates W542L and S621I double mutations in two ALS genes in maize"

### Additional file 1

#### Figure S1. Sequencing chromatograms from 7 prime-edited lines harboring W542L edits

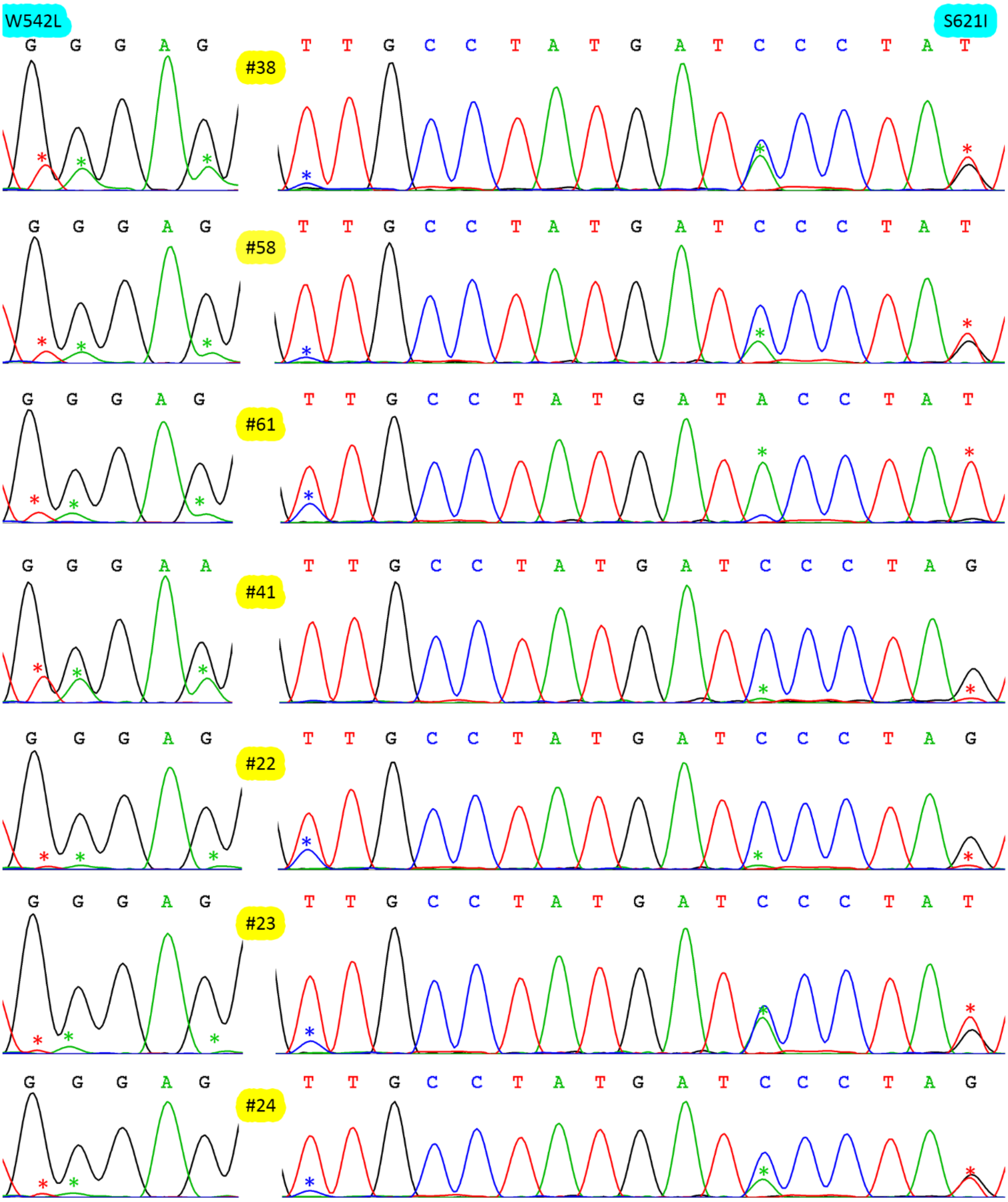

**Fig. S1. Sequencing chromatograms from 7 prime-edited lines harboring W542L edits.** Double peaks represent heterozygous or chimeric mutations and an asterisk indicates a mutation induced by PE. Note that the first asterisk of the S621I edits indicates the pegRNA scaffold-derived byproducts.

**Figure S2. Sequencing chromatograms from 2 prime-edited lines**

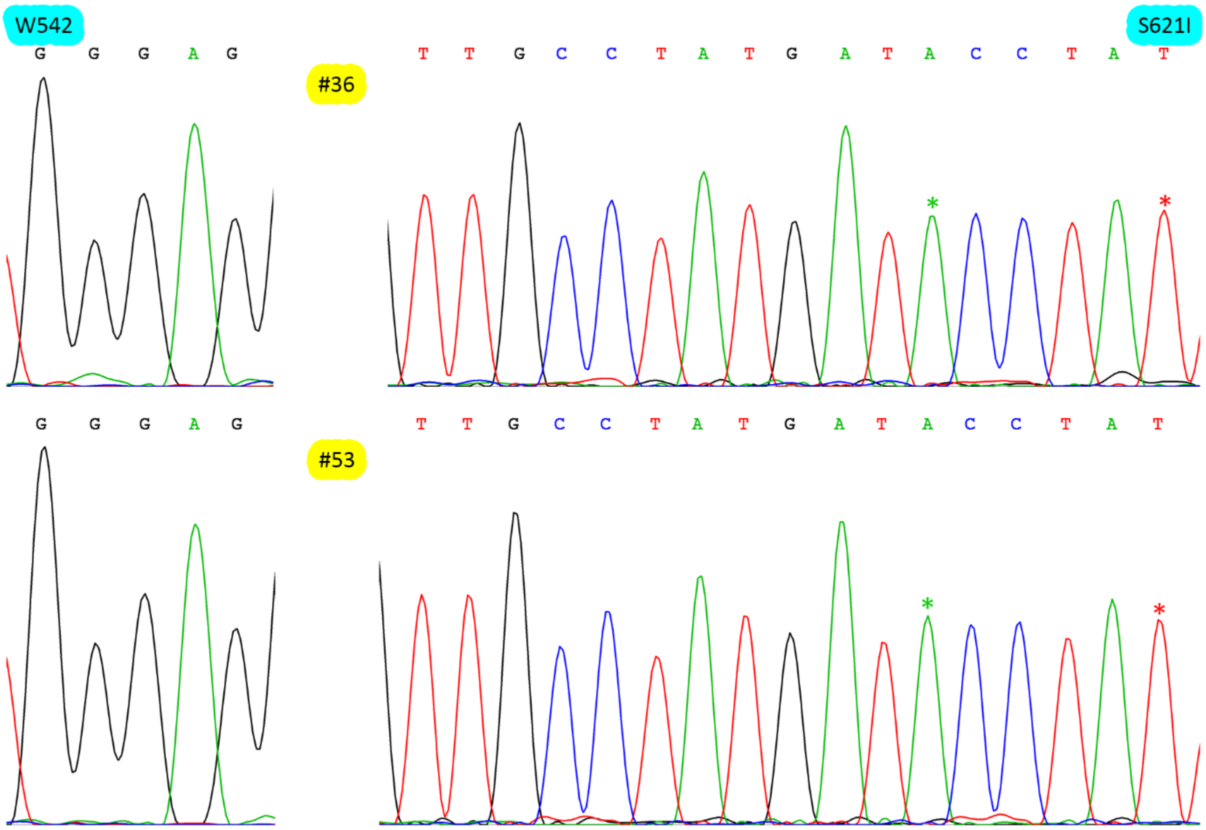

**Fig. S2. Sequencing chromatograms from 2 prime-edited lines harboring homozygous S621I edits.** An asterisk indicates a mutation induced by PE.

#### Figure S3. Prime-editing efficiencies in rice protoplasts for pegRNAs based on different expression strategies

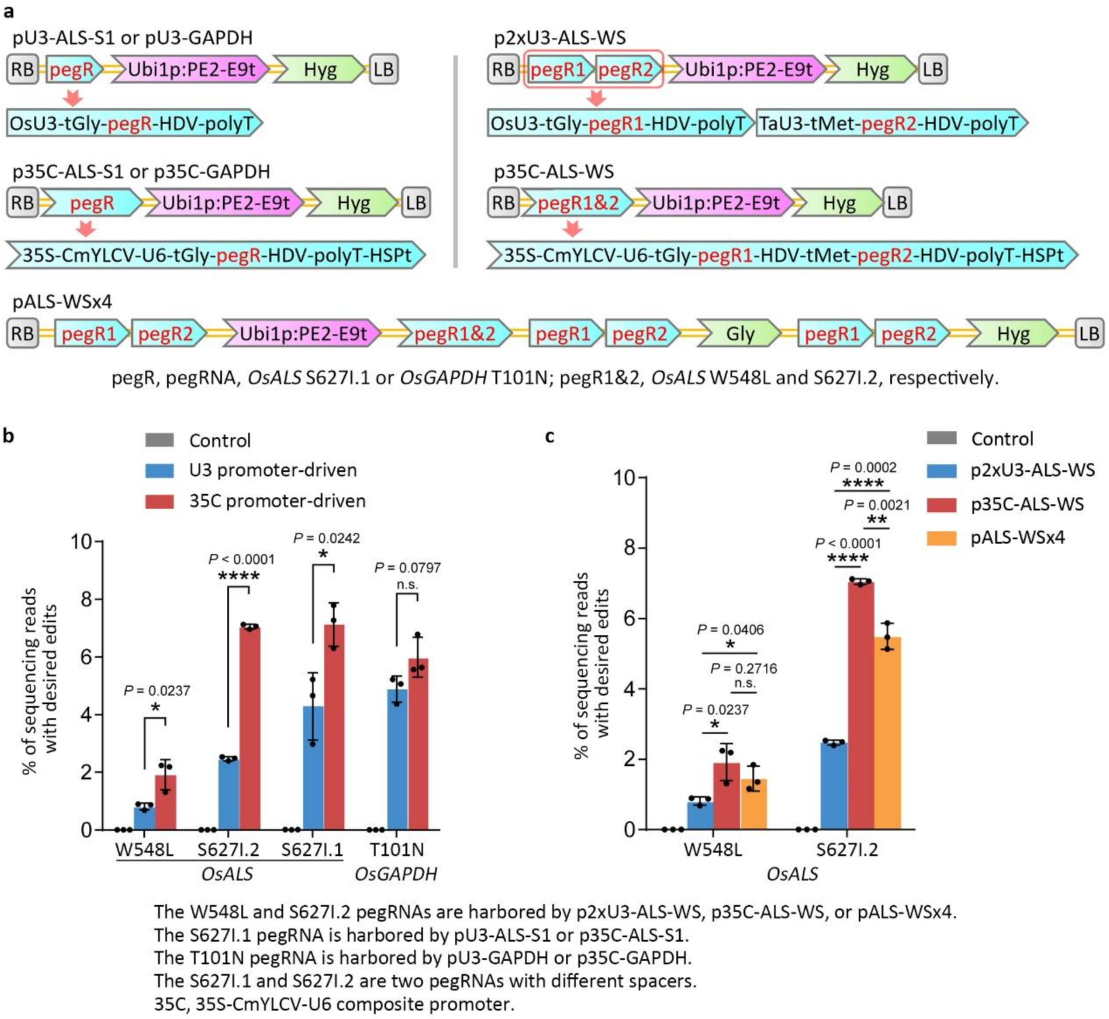

**Fig. S3. Prime-editing efficiencies in rice protoplasts for pegRNAs based on different expression strategies. a** T-DNA structures of the seven rice PE vectors. **b** Prime-editing efficiencies in rice protoplasts for pegRNAs driven by the 35S-CmYLCV-U6 composite promoter. An untreated protoplast sample served as a control. **c** Prime-editing efficiencies in rice protoplasts for pegRNAs based on the expression strategy of redoubling the number of expression cassettes. Efficiency (mean ± s.e.m.) was calculated from three independent experiments (n = 3). *P* values were obtained using the two-tailed Student's *t*-test. **P* < 0.05, ***P* < 0.01, ****P* < 0.001, *****P* < 0.0001. n.s., *P* > 0.05.

#### Table S1. Edits and byproducts revealed from cloned PCR fragments

**Table S1.** Edits and byproducts revealed from cloned PCR fragments

| Mutation | Line | Gene | Total | De | Sc | Re | To | De% | Sc% | Re% | To% | De/To | Sc/To | Re/To |
|---|---|---|---|---|---|---|---|---|---|---|---|---|---|---|
| **S621I** | **#3** | ***ALS1*** | 48 | 1 | 0 | 2 | 3 | 2.1% | 0 | 4.2% | 6.3% | 33.3% | 0 | 66.7% |
| | | ***ALS2*** | 43 | 3 | 0 | 0 | 3 | 7.0% | 0 | 0 | 7.0% | 100% | 0 | 0 |
| | | ***ALS1/2*** | 91 | 4 | 0 | 2 | 6 | 4.4% | 0 | 2.2% | 6.6% | 66.7% | 0 | 33.3% |
| | **#4** | ***ALS1*** | 41 | 41 | 0 | 0 | 41 | 100% | 0 | 0 | 100% | 100% | 0 | 0 |
| | | ***ALS2*** | 47 | 47 | 0 | 0 | 47 | 100% | 0 | 0 | 100% | 100% | 0 | 0 |
| | | ***ALS1/2*** | 88 | 88 | 0 | 0 | 88 | 100% | 0 | 0 | 100% | 100% | 0 | 0 |
| | **#5** | ***ALS1*** | 51 | 7 | 0 | 1 | 8 | 13.7% | 0 | 2.0% | 15.7% | 87.5% | 0 | 12.5% |
| | | ***ALS2*** | 27 | 5 | 0 | 1 | 6 | 18.5% | 0 | 3.7% | 22.2% | 83.3% | 0 | 16.7% |
| | | ***ALS1/2*** | 78 | 12 | 0 | 2 | 14 | 15.4% | 0 | 2.6% | 17.9% | 85.7% | 0 | 14.3% |
| | **#6** | ***ALS1*** | 40 | 3 | 1 | 0 | 4 | 7.5% | 2.5% | 0 | 10.0% | 75.0% | 25.0% | 0 |
| | | ***ALS2*** | 46 | 5 | 0 | 1 | 6 | 10.9% | 0 | 2.2% | 13.0% | 83.3% | 0 | 16.7% |
| | | ***ALS1/2*** | 86 | 8 | 1 | 1 | 10 | 9.3% | 1.2% | 1.2% | 11.6% | 80.0% | 10.0% | 10.0% |
| | **#8** | ***ALS1*** | 48 | 6 | 1 | 1 | 8 | 12.5% | 2.1% | 2.1% | 16.7% | 75.0% | 12.5% | 12.5% |
| | | ***ALS2*** | 45 | 2 | 4 | 1 | 7 | 4.4% | 8.9% | 2.2% | 15.6% | 28.6% | 57.1% | 14.3% |
| | | ***ALS1/2*** | 93 | 8 | 5 | 2 | 15 | 8.6% | 5.4% | 2.2% | 16.1% | 53.3% | 33.3% | 13.3% |
| | **#14** | ***ALS1*** | 45 | 7 | 0 | 1 | 8 | 15.6% | 0 | 2.2% | 17.8% | 87.5% | 0 | 12.5% |
| | | ***ALS2*** | 43 | 4 | 0 | 1 | 5 | 9.3% | 0 | 2.3% | 11.6% | 80.0% | 0 | 20.0% |
| | | ***ALS1/2*** | 88 | 11 | 0 | 2 | 13 | 12.5% | 0 | 2.3% | 14.8% | 84.6% | 0 | 15.4% |
| | **#15** | ***ALS1*** | 56 | 12 | 36 | 3 | 51 | 21.4% | 64.3% | 5.4% | 91.1% | 23.5% | 70.6% | 5.9% |
| | | ***ALS2*** | 21 | 6 | 4 | 2 | 12 | 28.6% | 19.0% | 9.5% | 57.1% | 50.0% | 33.3% | 16.7% |
| | | ***ALS1/2*** | 77 | 18 | 40 | 5 | 63 | 23.4% | 51.9% | 6.5% | 81.8% | 28.6% | 63.5% | 7.9% |
| | **#16** | ***ALS1*** | 54 | 3 | 4 | 2 | 9 | 5.6% | 7.4% | 3.7% | 16.7% | 33.3% | 44.4% | 22.2% |
| | | ***ALS2*** | 30 | 2 | 3 | 0 | 5 | 6.7% | 10.0% | 0 | 16.7% | 40.0% | 60.0% | 0 |
| | | ***ALS1/2*** | 84 | 5 | 7 | 2 | 14 | 6.0% | 8.3% | 2.4% | 16.7% | 35.7% | 50.0% | 14.3% |
| | **All-1** | ***ALS1*** | 383 | 80 | 42 | 10 | 132 | 20.9% | 11.0% | 2.6% | 34.5% | 60.6% | 31.8% | 7.6% |
| | | ***ALS2*** | 302 | 74 | 11 | 6 | 91 | 24.5% | 3.6% | 2.0% | 30.1% | 81.3% | 12.1% | 6.6% |
| | | ***ALS1/2*** | 685 | 154 | 53 | 16 | 223 | 22.5% | 7.7% | 2.3% | 32.6% | 69.1% | 23.8% | 7.2% |
| | **All-2** | ***ALS1*** | 286 | 27 | 6 | 7 | 40 | 9.4% | 2.1% | 2.4% | 14.0% | 67.5% | 15.0% | 17.5% |
| | | ***ALS2*** | 234 | 21 | 7 | 4 | 32 | 9.0% | 3.0% | 1.7% | 13.7% | 65.6% | 21.9% | 12.5% |
| | | ***ALS1/2*** | 520 | 48 | 13 | 11 | 72 | 9.2% | 2.5% | 2.1% | 13.8% | 66.7% | 18.1% | 15.3% |
| **W542L** | **#3** | ***ALS1*** | 48 | 0 | 0 | 0 | 0 | 0 | 0 | 0 | 0 | / | / | / |
| | | ***ALS2*** | 43 | 0 | 0 | 0 | 0 | 0 | 0 | 0 | 0 | / | / | / |
| | | ***ALS1/2*** | 91 | 0 | 0 | 0 | 0 | 0 | 0 | 0 | 0 | / | / | / |
| | **#4** | ***ALS1*** | 41 | 0 | 0 | 0 | 0 | 0 | 0 | 0 | 0 | / | / | / |
| | | ***ALS2*** | 47 | 0 | 0 | 0 | 0 | 0 | 0 | 0 | 0 | / | / | / |
| | | ***ALS1/2*** | 88 | 0 | 0 | 0 | 0 | 0 | 0 | 0 | 0 | / | / | / |
| | **#5** | ***ALS1*** | 51 | 2 | 0 | 0 | 2 | 3.9% | 0 | 0 | 3.9% | 100% | 0 | 0 |
| | | ***ALS2*** | 27 | 1 | 0 | 0 | 1 | 3.7% | 0 | 0 | 3.7% | 100% | 0 | 0 |
| | | ***ALS1/2*** | 78 | 3 | 0 | 0 | 3 | 3.8% | 0 | 0 | 3.8% | 100% | 0 | 0 |
| | **#6** | ***ALS1*** | 40 | 0 | 0 | 1 | 1 | 0 | 0 | 2.5% | 2.5% | 0 | 0 | 100% |
| | | ***ALS2*** | 46 | 1 | 0 | 0 | 1 | 2.2% | 0 | 0 | 2.2% | 100% | 0 | 0 |
| | | ***ALS1/2*** | 86 | 1 | 0 | 1 | 2 | 1.2% | 0 | 1.2% | 2.3% | 50.0% | 0 | 50.0% |
| | **#8** | ***ALS1*** | 48 | 0 | 0 | 0 | 0 | 0 | 0 | 0 | 0 | / | / | / |
| | | ***ALS2*** | 45 | 0 | 0 | 0 | 0 | 0 | 0 | 0 | 0 | / | / | / |
| | | ***ALS1/2*** | 93 | 0 | 0 | 0 | 0 | 0 | 0 | 0 | 0 | / | / | / |
| | **#14** | ***ALS1*** | 45 | 0 | 0 | 0 | 0 | 0 | 0 | 0 | 0 | / | / | / |
| | | ***ALS2*** | 43 | 0 | 0 | 0 | 0 | 0 | 0 | 0 | 0 | / | / | / |
| | | ***ALS1/2*** | 88 | 0 | 0 | 0 | 0 | 0 | 0 | 0 | 0 | / | / | / |
| | **#15** | ***ALS1*** | 56 | 1 | 0 | 0 | 1 | 1.8% | 0 | 0 | 1.8% | 100% | 0 | 0 |
| | | ***ALS2*** | 21 | 0 | 0 | 0 | 0 | 0 | 0 | 0 | 0 | / | / | / |

| | | | | | | | | | | | | | | |
|---|---|---|---|---|---|---|---|---|---|---|---|---|---|---|
| | | ***ALS1/2*** | 77 | 1 | 0 | 0 | 1 | 1.3% | 0 | 0 | 1.3% | 100% | 0 | 0 |
| | **#16** | ***ALS1*** | 54 | 1 | 1 | 0 | 2 | 1.9% | 1.9% | 0 | 3.7% | 50.0% | 50 | 0 |
| | | ***ALS2*** | 30 | 1 | 0 | 0 | 1 | 3.3% | 0 | 0 | 3.3% | 100% | 0 | 0 |
| | | ***ALS1/2*** | 84 | 2 | 1 | 0 | 3 | 2.4% | 1.2% | 0 | 3.6% | 66.7% | 33.3% | 0 |
| | **All-1** | ***ALS1*** | 383 | 4 | 1 | 1 | 6 | 1.0% | 0.3% | 0.3% | 1.6% | 66.7% | 16.7% | 16.7% |
| | | ***ALS2*** | 302 | 3 | 0 | 0 | 3 | 1.0% | 0 | 0 | 1.0% | 100% | 0 | 0 |
| | | ***ALS1/2*** | 685 | 7 | 1 | 1 | 9 | 1.0% | 0.1% | 0.1% | 1.3% | 77.8% | 11.1% | 11.1% |
| **P165S (PE3b)** | **#51** | ***ALS1*** | 46 | 0 | 0 | 0 | 0 | 0 | 0 | 0 | 0 | / | / | / |
| | | ***ALS2*** | 53 | 0 | 0 | 0 | 0 | 0 | 0 | 0 | 0 | / | / | / |
| | | ***ALS1/2*** | 99 | 0 | 0 | 0 | 0 | 0 | 0 | 0 | 0 | / | / | / |
| | **#161** | ***ALS1*** | 36 | 0 | 0 | 0 | 0 | 0 | 0 | 0 | 0 | / | / | / |
| | | ***ALS2*** | 25 | 0 | 0 | 0 | 0 | 0 | 0 | 0 | 0 | / | / | / |
| | | ***ALS1/2*** | 61 | 0 | 0 | 0 | 0 | 0 | 0 | 0 | 0 | / | / | / |
| | **#162** | ***ALS1*** | 26 | 0 | 0 | 0 | 0 | 0 | 0 | 0 | 0 | / | / | / |
| | | ***ALS2*** | 28 | 0 | 0 | 0 | 0 | 0 | 0 | 0 | 0 | / | / | / |
| | | ***ALS1/2*** | 54 | 0 | 0 | 0 | 0 | 0 | 0 | 0 | 0 | / | / | / |
| | **All-1** | ***ALS1*** | 108 | 0 | 0 | 0 | 0 | 0 | 0 | 0 | 0 | / | / | / |
| | | ***ALS2*** | 106 | 0 | 0 | 0 | 0 | 0 | 0 | 0 | 0 | / | / | / |
| | | ***ALS1/2*** | 214 | 0 | 0 | 0 | 0 | 0 | 0 | 0 | 0 | / | / | / |
| **P165S (PE3)** | **#54** | ***ALS1*** | 33 | 0 | 0 | 0 | 0 | 0 | 0 | 0 | 0 | / | / | / |
| | | ***ALS2*** | 31 | 0 | 0 | 0 | 0 | 0 | 0 | 0 | 0 | / | / | / |
| | | ***ALS1/2*** | 64 | 0 | 0 | 0 | 0 | 0 | 0 | 0 | 0 | / | / | / |
| | **#55** | ***ALS1*** | 31 | 0 | 0 | 0 | 0 | 0 | 0 | 0 | 0 | / | / | / |
| | | ***ALS2*** | 25 | 0 | 0 | 0 | 0 | 0 | 0 | 0 | 0 | / | / | / |
| | | ***ALS1/2*** | 56 | 0 | 0 | 0 | 0 | 0 | 0 | 0 | 0 | / | / | / |
| | **#56** | ***ALS1*** | 41 | 0 | 0 | 0 | 0 | 0 | 0 | 0 | 0 | / | / | / |
| | | ***ALS2*** | 34 | 0 | 0 | 0 | 0 | 0 | 0 | 0 | 0 | / | / | / |
| | | ***ALS1/2*** | 75 | 0 | 0 | 0 | 0 | 0 | 0 | 0 | 0 | / | / | / |
| | **All-1** | ***ALS1*** | 105 | 0 | 0 | 0 | 0 | 0 | 0 | 0 | 0 | / | / | / |
| | | ***ALS2*** | 90 | 0 | 0 | 0 | 0 | 0 | 0 | 0 | 0 | / | / | / |
| | | ***ALS1/2*** | 195 | 0 | 0 | 0 | 0 | 0 | 0 | 0 | 0 | / | / | / |

Total, total number of sequenced clones; De, Sc, Re, or To, No. of clones harboring specified mutations; De, desired edits; Sc, pegRNA scaffold-derived byproducts; Re, double-strand even DNA repair-derived byproducts; To, total No. of cloned fragments harboring all the three types of mutations; De%, Sc%, Re%, or To%, ratio of clones harboring specified mutations to total number of sequenced clones; All-1, value for all the lines; All-2, value for all the lines but #4 and #15.

#### Table S2. Analysis of mutations in T0 transgenic plants by NGS with 0.5% threshold

**Table S2.** Analysis of mutations in T0 transgenic plants by NGS with 0.5% threshold

| Mutation | Line | Gene | De% | Sc% | Re% | To% | De/To | Sc/To | Re/To |
|---|---|---|---|---|---|---|---|---|---|
| **S621I** | **#1** | ***ALS1*** | 4.2% | 0.9% | 0.9% | 6.0% | 70.8% | 14.9% | 14.4% |
| | | ***ALS2*** | 4.0% | 0.9% | 0.8% | 5.7% | 69.4% | 15.8% | 14.7% |
| | | ***ALS1/2*** | 4.1% | 0.9% | 0.8% | 5.8% | 70.1% | 15.3% | 14.5% |
| | **#3** | ***ALS1*** | 3.7% | 0.7% | 0.7% | 5.1% | 72.9% | 13.4% | 13.8% |
| | | ***ALS2*** | 3.6% | 0.8% | 0.7% | 5.1% | 70.6% | 16.0% | 13.4% |
| | | ***ALS1/2*** | 3.7% | 0.7% | 0.7% | 5.1% | 71.7% | 14.7% | 13.6% |
| | **#4** | ***ALS1*** | 100% | 0 | 0 | 100% | 100% | 0 | 0 |
| | | ***ALS2*** | 100% | 0 | 0 | 100% | 100% | 0 | 0 |
| | | ***ALS1/2*** | 100% | 0 | 0 | 100% | 100% | 0 | 0 |
| | **#5** | ***ALS1*** | 9.4% | 1.6% | 1.8% | 12.7% | 73.8% | 12.3% | 13.9% |
| | | ***ALS2*** | 9.4% | 1.6% | 1.8% | 12.8% | 73.4% | 12.4% | 14.2% |
| | | ***ALS1/2*** | 9.4% | 1.6% | 1.8% | 12.8% | 73.6% | 12.3% | 14.1% |
| | **#6** | ***ALS1*** | 8.0% | 1.3% | 2.3% | 11.6% | 68.9% | 11.3% | 19.8% |
| | | ***ALS2*** | 7.7% | 1.7% | 2.0% | 11.4% | 67.9% | 14.6% | 17.5% |
| | | ***ALS1/2*** | 7.9% | 1.5% | 2.1% | 11.5% | 68.4% | 12.9% | 18.7% |
| | **#8** | ***ALS1*** | 12.6% | 3.2% | 2.6% | 18.5% | 68.3% | 17.4% | 14.3% |
| | | ***ALS2*** | 12.4% | 3.2% | 2.9% | 18.5% | 67.3% | 17.3% | 15.5% |
| | | ***ALS1/2*** | 12.5% | 3.2% | 2.7% | 18.5% | 67.8% | 17.3% | 14.9% |
| | **#9** | ***ALS1*** | 4.0% | 0.8% | 1.2% | 6.0% | 66.9% | 12.8% | 20.3% |
| | | ***ALS2*** | 3.7% | 0.9% | 1.0% | 5.6% | 66.2% | 16.1% | 17.7% |
| | | ***ALS1/2*** | 3.9% | 0.8% | 1.1% | 5.8% | 66.6% | 14.3% | 19.1% |
| | **#12** | ***ALS1*** | 1.6% | 0 | 0 | 1.6% | 100% | 0 | 0 |
| | | ***ALS2*** | 1.7% | 0 | 0 | 1.7% | 100% | 0 | 0 |
| | | ***ALS1/2*** | 1.7% | 0 | 0 | 1.7% | 100% | 0 | 0 |
| | **#13** | ***ALS1*** | 3.1% | 0 | 0 | 3.1% | 100% | 0 | 0 |
| | | ***ALS2*** | 2.8% | 0 | 0 | 2.8% | 100% | 0 | 0 |
| | | ***ALS1/2*** | 2.9% | 0 | 0 | 2.9% | 100% | 0 | 0 |
| | **#14** | ***ALS1*** | 14.5% | 0.6% | 4.8% | 19.9% | 72.9% | 2.9% | 24.2% |
| | | ***ALS2*** | 15.1% | 0.7% | 2.8% | 18.6% | 81.4% | 3.5% | 15.1% |
| | | ***ALS1/2*** | 14.8% | 0.6% | 3.8% | 19.2% | 77.0% | 3.2% | 19.8% |
| | **#15** | ***ALS1*** | 21.3% | 56.4% | 5.3% | 83.0% | 25.7% | 68.0% | 6.3% |
| | | ***ALS2*** | 39.5% | 15.1% | 8.8% | 63.4% | 62.3% | 23.9% | 13.8% |
| | | ***ALS1/2*** | 30.4% | 35.8% | 7.0% | 73.2% | 41.6% | 48.9% | 9.6% |
| | **#16** | ***ALS1*** | 11.6% | 3.0% | 3.3% | 17.8% | 64.8% | 16.8% | 18.4% |
| | | ***ALS2*** | 13.0% | 2.9% | 3.4% | 19.3% | 67.2% | 15.2% | 17.6% |
| | | ***ALS1/2*** | 12.3% | 3.0% | 3.3% | 18.6% | 66.1% | 16.0% | 17.9% |
| | **Aver-1** | ***ALS1*** | 16.2% | 5.7% | 1.9% | 23.8% | 68.0% | 24.0% | 8.0% |
| | | ***ALS2*** | 17.7% | 2.3% | 2.0% | 22.1% | 80.4% | 10.5% | 9.1% |
| | | ***ALS1/2*** | 17.0% | 4.0% | 2.0% | 22.9% | 74.0% | 17.5% | 8.5% |
| | **Aver-2** | ***ALS1*** | 7.3% | 1.2% | 1.8% | 10.2% | 71.1% | 11.7% | 17.2% |
| | | ***ALS2*** | 7.3% | 1.3% | 1.5% | 10.2% | 72.4% | 12.5% | 15.2% |

| | | | | | | | | | |
|---|---|---|---|---|---|---|---|---|---|
| | | ***ALS1/2*** | 7.3% | 1.2% | 1.6% | 10.2% | 71.7% | 12.1% | 16.2% |
| **W542L** | **#5** | ***ALS1*** | 1.3% | 0 | 0 | 1.3% | 100% | 0 | 0 |
| | | ***ALS2*** | 1.0% | 0 | 0 | 1.0% | 100% | 0 | 0 |
| | | ***ALS1/2*** | 1.2% | 0 | 0 | 1.2% | 100% | 0 | 0 |
| | **#6** | ***ALS1*** | 0.8% | 0 | 0 | 0.8% | 100% | 0 | 0 |
| | | ***ALS2*** | 0.5% | 0 | 0 | 0.5% | 100% | 0 | 0 |
| | | ***ALS1/2*** | 0.7% | 0 | 0 | 0.7% | 100% | 0 | 0 |
| | **#8** | ***ALS1*** | 1.0% | 0 | 0 | 1.0% | 100% | 0 | 0 |
| | | ***ALS2*** | 0.9% | 0 | 0 | 0.9% | 100% | 0 | 0 |
| | | ***ALS1/2*** | 0.9% | 0 | 0 | 0.9% | 100% | 0 | 0 |
| | **#13** | ***ALS1*** | 1.3% | 0 | 0 | 1.3% | 100% | 0 | 0 |
| | | ***ALS2*** | 0.6% | 0 | 0 | 0.6% | 100% | 0 | 0 |
| | | ***ALS1/2*** | 0.9% | 0 | 0 | 0.9% | 100% | 0 | 0 |
| | **#15** | ***ALS1*** | 4.6% | 0 | 0 | 4.6% | 100% | 0 | 0 |
| | | ***ALS2*** | 2.9% | 0 | 0 | 2.9% | 100% | 0 | 0 |
| | | ***ALS1/2*** | 3.8% | 0 | 0 | 3.8% | 100% | 0 | 0 |
| | **#16** | ***ALS1*** | 1.6% | 0 | 0 | 1.6% | 100% | 0 | 0 |
| | | ***ALS2*** | 1.6% | 0 | 0 | 1.6% | 100% | 0 | 0 |
| | | ***ALS1/2*** | 1.6% | 0 | 0 | 1.6% | 100% | 0 | 0 |
| | **Aver-1** | ***ALS1*** | 1.8% | 0 | 0 | 1.8% | 100% | 0 | 0 |
| | | ***ALS2*** | 1.2% | 0 | 0 | 1.2% | 100% | 0 | 0 |
| | | ***ALS1/2*** | 1.5% | 0 | 0 | 1.5% | 100% | 0 | 0 |

De, Sc, Re, or To, No. of clones harboring specified mutations; De, desired edits; Sc, pegRNA scaffold-derived byproducts; Re, double-strand even DNA repair-derived byproducts; To, total No. of cloned fragments harboring all the three types of mutations; De%, Sc%, Re%, or To%, ratio of clones harboring specified mutations to total number of NGS reads; Aver-1, average value for all the lines; Aver-2, average value for all the lines but #4 and #15.

#### Table S3. Edits and byproducts from the 4 additional lines

**Table S3.** Edits and byproducts from the 4 additional lines

| Mutation | Line | Gene | Total | De | Sc | Re | To | De% | Sc% | Re% | To% | De/To | Sc/To | Re/To |
|---|---|---|---|---|---|---|---|---|---|---|---|---|---|---|
| S621I | #38 | *ALS1* | 52 | 12 | 3 | 2 | 17 | 23.1% | 5.8% | 3.8% | 32.7% | 70.6% | 17.6% | 11.8% |
| | | *ALS2* | 61 | 25 | 4 | 2 | 31 | 41.0% | 6.6% | 3.3% | 50.8% | 80.6% | 12.9% | 6.5% |
| | | *ALS1/2* | 113 | 37 | 7 | 4 | 48 | 32.7% | 6.2% | 3.5% | 42.5% | 77.1% | 14.6% | 8.3% |
| | #41 | *ALS1* | 62 | 4 | 1 | 1 | 6 | 6.5% | 1.6% | 1.6% | 9.7% | 66.7% | 16.7% | 16.7% |
| | | *ALS2* | 51 | 5 | 2 | 0 | 7 | 9.8% | 3.9% | 0 | 13.7% | 71.4% | 28.6% | 0 |
| | | *ALS1/2* | 113 | 9 | 3 | 1 | 13 | 8.0% | 2.7% | 0.9% | 11.5% | 69.2% | 23.1% | 7.7% |
| | #58 | *ALS1* | 47 | 6 | 1 | 14 | 21 | 12.8% | 2.1% | 29.8% | 44.7% | 28.6% | 4.8% | 66.7% |
| | | *ALS2* | 45 | 5 | 4 | 6 | 15 | 11.1% | 8.9% | 13.3% | 33.3% | 33.3% | 26.7% | 40.0% |
| | | *ALS1/2* | 92 | 11 | 5 | 20 | 36 | 12.0% | 5.4% | 21.7% | 39.1% | 30.6% | 13.9% | 55.6% |
| | #61 | *ALS1* | 49 | 47 | 2 | 0 | 49 | 95.9% | 4.1% | 0 | 100.0% | 95.9% | 4.1% | 0 |
| | | *ALS2* | 57 | 13 | 25 | 1 | 39 | 22.8% | 43.9% | 1.8% | 68.4% | 33.3% | 64.1% | 2.6% |
| | | *ALS1/2* | 106 | 60 | 27 | 1 | 88 | 56.6% | 25.5% | 0.9% | 83.0% | 68.2% | 30.7% | 1.1% |
| | All | *ALS1* | 210 | 69 | 7 | 17 | 93 | 32.9% | 3.3% | 8.1% | 44.3% | 74.2% | 7.5% | 18.3% |
| | | *ALS2* | 214 | 48 | 35 | 9 | 92 | 22.4% | 16.4% | 4.2% | 43.0% | 52.2% | 38.0% | 9.8% |
| | | *ALS1/2* | 424 | 117 | 42 | 26 | 185 | 27.6% | 9.9% | 6.1% | 43.6% | 63.2% | 22.7% | 14.1% |
| W542L | #38 | *ALS1* | 52 | 4 | 0 | 0 | 4 | 7.7% | 0 | 0 | 7.7% | 100.0% | 0 | 0 |
| | | *ALS2* | 61 | 33 | 0 | 1 | 34 | 54.1% | 0 | 1.6% | 55.7% | 97.1% | 0.0% | 2.9% |
| | | *ALS1/2* | 113 | 37 | 0 | 1 | 38 | 32.7% | 0 | 0.9% | 33.6% | 97.4% | 0.0% | 2.6% |
| | #41 | *ALS1* | 62 | 0 | 0 | 0 | 0 | 0 | 0 | 0 | 0 | / | / | / |
| | | *ALS2* | 51 | 31 | 0 | 0 | 31 | 60.8% | 0 | 0 | 60.8% | 100.0% | 0 | 0 |
| | | *ALS1/2* | 113 | 31 | 0 | 0 | 31 | 27.4% | 0 | 0 | 27.4% | 100.0% | 0 | 0 |
| | #58 | *ALS1* | 47 | 7 | 0 | 0 | 7 | 14.9% | 0 | 0 | 14.9% | 100.0% | 0 | 0 |
| | | *ALS2* | 45 | 6 | 0 | 1 | 7 | 13.3% | 0 | 2.2% | 15.6% | 85.7% | 0 | 14.3% |
| | | *ALS1/2* | 92 | 13 | 0 | 1 | 14 | 14.1% | 0 | 1.1% | 15.2% | 92.9% | 0 | 7.1% |
| | #61 | *ALS1* | 49 | 3 | 0 | 1 | 4 | 6.1% | 0 | 2.0% | 8.2% | 75.0% | 0 | 25.0% |
| | | *ALS2* | 57 | 9 | 0 | 4 | 13 | 15.8% | 0 | 7.0% | 22.8% | 69.2% | 0 | 30.8% |
| | | *ALS1/2* | 106 | 12 | 0 | 5 | 17 | 11.3% | 0 | 4.7% | 16.0% | 70.6% | 0 | 29.4% |
| | All | *ALS1* | 210 | 14 | 0 | 1 | 15 | 6.7% | 0 | 0.5% | 7.1% | 93.3% | 0 | 6.7% |
| | | *ALS2* | 214 | 79 | 0 | 6 | 85 | 36.9% | 0 | 2.8% | 39.7% | 92.9% | 0 | 7.1% |
| | | *ALS1/2* | 424 | 93 | 0 | 7 | 100 | 21.9% | 0 | 1.7% | 23.6% | 93.0% | 0 | 7.0% |

Total, total number of sequenced clones; De, Sc, Re, or To, No. of clones harboring specified mutations; De, desired edits; Sc, pegRNA scaffold-derived byproducts; Re, double-strand even DNA repair-derived byproducts; To, total No. of cloned fragments harboring all the three types of mutations; De%, Sc%, Re%, or To%, ratio of clones harboring specified mutations to total number of sequenced clones; All, value for all the lines.

### Table S4. Prime-editing efficiency in rice protoplasts analyzed by NGS

**Table S4.** Prime-editing efficiency in rice protoplasts analyzed by NGS

| pegRNA/Mutation | Vector | Promoter | Sequencing reads with desired edits (%) | | | |
|---|---|---|---|---|---|---|
| | | | n=1 | n=2 | n=3 | Aver. |
| W548L | / | / | 0.0 | 0.0 | 0.0 | 0.0 |
| | p2xU3-ALS-WS | U3 | 0.7 | 0.9 | 0.9 | 0.8 |
| | p35C-ALS-WS | 35C | 2.2 | 1.3 | 2.2 | 1.9 |
| | pALS-WSx4 | U3&35C | 1.8 | 1.2 | 1.4 | 1.5 |
| S627I.2 | / | / | 0.0 | 0.0 | 0.0 | 0.0 |
| | p2xU3-ALS-WS | U3 | 2.5 | 2.4 | 2.5 | 2.5 |
| | p35C-ALS-WS | 35C | 7.1 | 7.0 | 7.1 | 7.0 |
| | pALS-WSx4 | U3&35C | 5.9 | 5.1 | 5.5 | 5.5 |
| S627I.1 | / | / | 0.0 | 0.0 | 0.0 | 0.0 |
| | pU3-ALS-S1 | U3 | 3.0 | 5.2 | 4.7 | 4.3 |
| | p35C-ALS-S1 | 35C | 7.3 | 6.3 | 7.8 | 7.1 |
| T101N | / | / | 0.0 | 0.0 | 0.0 | 0.0 |
| | pU3-GAPDH | U3 | 5.1 | 4.4 | 5.2 | 4.9 |
| | p35C-GAPDH | 35C | 5.6 | 5.6 | 6.8 | 6.0 |

35C, CaMV35S-CmYLCV-U6 composite promoter. Aver., average value for three independent experiments (n = 1, 2, 3).

#### Table S5. Sequences of primers, targets, and rtT-PBS of pegRNAs

**Table S5.** Sequences of primers, targets, and rtT-PBS of pegRNAs

| Sequence name | Sequence | Purpose |
|---|---|---|
| oHEASmE-F | AGCTTGCTGAATTCGTCAACACCTGCAACACTAGT | Vector construction |
| oHEASmE-R | AATTACTAGTGTTGCAGGTGTTGACGAATTCAGCA | |
| OsU3p-AsF | ATTTATTTAGGCGCGCCAGTAATTCATCCAGGTCAC | |
| TaU3t-EcR | AACACCATGAATTCAAGATGTTGTACTTCTGAA | |
| oiSce-HSF | AGCTTGAGACCATTACCCTGTTATCCCTAGGTCTCGAATTA | |
| oiSce-HSR | CTAGTAATTCGAGACCTAGGGATAACAGGGTAATGGTCTCA | |
| ALS1&2P-F | CGTCATCGCCAACCACCTCTTC | Analysis of the P165S mutation by Sanger sequencing |
| ALS1&2P-R | CCATCTGCTGCTGGATGTCCTTG | |
| ALS1&2WS-F | CTTGGGGCTATGGGATTTGGTTTGC | Analysis of the W542L and S621I mutations by Sanger sequencing |
| ALS1&2WS-R | TACACAGTCCTGCCATCACCATC | |
| ALS1&2WS-F2 | CTTCTGTGGCCAACCCAGGTGT | |
| ALS1&2P-NGSF | GGAGTGAGTACGGTGTGCCGTCTGCATCGCCACCTC | Analysis of the P165S mutation by NGS |
| ALS1&2P-NGSR | GAGTTGGATGCTGGATGGACGATGGGCGTCTCCTG | |
| ALS1&2W-NGSF | GGAGTGAGTACGGTGTGCGATCCGAATTGAGAACCTCC | Analysis of the W542L mutation by NGS |
| ALS1&2W-NGSR | GAGTTGGATGCTGGATGGCATTCTCTGGGTTTCCCAAG | |
| ALS1&2S-NGSF | GGAGTGAGTACGGTGTGCAGGGCCGTACCTCTTGGATA | Analysis of the S621I mutation by NGS |
| ALS1&2S-NGSR | GAGTTGGATGCTGGATGGACAGTCCTGCCATCACCAT | |
| P35C-GAPDH-T101N/F | AGTTCCGGTAGCGAGCGTGGAACTGATG | Analysis of the T101N mutation of *OsGAPDH* by NGS |
| p35C-GAPDH-T101N/R | AGTCAACTTGCTTGTATGCAATCCCATGGG | |
| pU3-GAPDH-T101N/F | CCGTCCGGTAGCGAGCGTGGAACTGATG | |
| pU3-GAPDH-T101N/R | ATGTCACTTGCTTGTATGCAATCCCATGGG | |
| GAPDH-Control/F | CGTACGGGTAGCGAGCGTGGAACTGATG | |
| GAPDH-Control/R | GTTTCGCTTGCTTGTATGCAATCCCATGGG | |
| p2xU3-ALS-WS-W548L/F | CGATGTTTCAGGAGCTGGCATTGATC | Analysis of the W548L mutation of *OsALS* by NGS |
| p2xU3-ALS-WS-W548L/R | TGACCAAGCAATAGTCACAAAATCTGG | |
| p35C-ALS-WS-W548L/F | ACAGTGTTCAGGAGCTGGCATTGATC | |
| p35C-ALS-WS-W548L/R | GCCAATAGCAATAGTCACAAAATCTGG | |
| pALS-WSx4-W548L/F | CAGATCTTCAGGAGCTGGCATTGATC | |
| pALS-WSx4-W548L/R | CTTGTAAGCAATAGTCACAAAATCTGG | |
| ALS-W548L-Control/F | TAGCTTTTCAGGAGCTGGCATTGATC | |
| ALS-W548L-Control/R | GGCTACAGCAATAGTCACAAAATCTGG | |
| p2xU3-ALS-WS-S627I.2/F | CGATGTCCGCCATCAAGAAGATGC | Analysis of the S627I mutation of *OsALS* by NGS |
| p2xU3-ALS-WS-S627I.2/R | TGACCATTCAGGTCAAACATAGGCCG | |
| p35C-ALS-WS-S627I.2/F | ACAGTGCCGCCATCAAGAAGATGC | |
| p35C-ALS-WS-S627I.2/R | GCCAATTTCAGGTCAAACATAGGCCG | |
| pALS-WSx4-S627I.2/F | CAGATCCCGCCATCAAGAAGATGC | |
| pALS-WSx4-S627I.2/R | CTTGTATTCAGGTCAAACATAGGCCG | |
| p35C-ALS-S627I.1/F | ATCACGCCGCCATCAAGAAGATGC | |
| p35C-ALS-S627I.1/R | TTAGGCTTCAGGTCAAACATAGGCCG | |
| pU3-ALS-S627I.1/F | ACTTGACCGCCATCAAGAAGATGC | |
| pU3-ALS-S627I.1/R | GATCAGTTCAGGTCAAACATAGGCCG | |

| | | |
|---|---|---|
| ALS-S627I-Control/F | TAGCTTCCGCCATCAAGAAGATGC | |
| ALS-S627I-Control/R | GGCTACTTCAGGTCAAACATAGGCCG | |
| ALS1&2P-T1 | TCGGTGCCAATCATGCGTCGCGG | Two pegRNAs and two sgRNAs for the P165S mutation |
| P165S-rtT/PBS | GGACAGGTGAGTCGA/CGCATGATTG | |
| ALS1&2P-T2 | CAGGTGAGTCGACGCATGATTGG | |
| P165S-rtT/PBS2 | GGACAGGTGAGTCGA/CGCATGATTGGCA | |
| ALS1&2P-T2b | YCAGGAGACGCCCATCGTCGAGG (Y = T or C) | |
| ALS1&2W-T1 | SGGGATGGTGGTGCAGTGGGAGG (S = C or G) | One pegRNA and one sgRNA for the W542L mutation |
| W542L-rtT/PBS | TAGAACCTGTCTTCTA/ACTGCACCACCAT | |
| ALS1W-T2 | CTAACTGCACCACCATCCCGAGG | |
| ALS1&2S-T1 | CCTTGAAAGCCCCACCACTAGGG | One pegRNA and one sgRNA for the S621I mutation |
| S621I-rtT/PBS | TGCCTATGATACCTAT/TGGTGGGGCTTTC | |
| ALS1&2S-T2 | GATATGATCCTGGATGGTGATGG | |
| OsALS-S627I.1-T1 | GTGCTGCCTATGATCCCAAGTGG | One pegRNA for the *OsALS* S627I mutation |
| OsALS-S627I.1-rtT/PBS | TGAATGCGCCCCCAaTT/GGGATCATAG | |
| OsGAPDH-T1 | GAGTATGTCGTGGAGTCCACCGG | One pegRNA for the *OsGAPDH* T101N mutation |
| OsGAPDH-rtT/PBS | AGTGAAGACACCGtTG/GACTCCACGACA | |
| OsALS-W548L-T1 | GGGTATGGTGGTGCAATGGGAGG | One pegRNA for the *OsALS* W548L mutation |
| OsALS-W548L-rtT/PBS | AAACCTATCtTCta/ATTGCACCACCAT | |
| OsALS-S627I.2-T1 | CCTTGAATGCGCCCCCACTTGGG | Another pegRNA for the *OsALS* S627I mutation |
| OsALS-S627I.2-rtT/PBS | TGCCTATGATaCCAAt/TGGGGGCGCATTC | |

#### Supplemental material. Sequences of PE2 and pegRNA expression cassettes

##### Maize codon-optimized PE2

NLS-SpCas9H840A-linker-M_MLV_RT-NLS

atgaagaggacagccgatggcagcgagttcgagagccctaagaagaagaggaaggtggacaagaagtactcgatcggcctcgatattgggactaactctgttggct
gggccgtgatcaccgacgagtacaaggtgccctcaaagaagttcaaggtcctgggcaacaccgatcggcattccatcaagaagaatctcattggcgctctcctgttcg
acagcggcgagacggctgaggctacgcggctcaagcgcaccgcccgcaggcggtacacgcgcaggaagaatcgcatctgctacctgcaggagattttctccaacga
gatggcgaaggttgacgattctttcttccacaggctggaggagtcattcctcgtggaggaggataagaagcacgagcggcatccaatcttcggcaacattgtcgacga
ggttgcctaccacgagaagtaccctacgatctaccatctgcggaagaagctcgtggactccacagataaggcggacctccgcctgatctacctcgctctggcccacat
gattaagttcaggggccatttcctgatcgagggggatctcaacccggacaatagcgatgttgacaagctgttcatccagctcgtgcagacgtacaaccagctcttcgag
gagaaccccattaatgcgtcaggcgtcgacgcgaaggctatcctgtccgctaggctctcgaagtctcggcgcctcgagaacctgatcgcccagctgccgggcgagaag
aagaacggcctgttcgggaatctcattgcgctcagcctggggctcacgcccaacttcaagtcgaatttcgatctcgctgaggacgccaagctgcagctctccaaggac
acatacgacgatgacctggataacctcctggcccagatcggcgatcagtacgcggacctgttcctcgctgccaagaatctgtcggacgccatcctcctgtctgatattct
cagggtgaacaccgagattacgaaggctccgctctcagcctccatgatcaagcgctacgacgagcaccatcaggatctgaccctcctgaaggcgctggtcaggcagc
agctccccgagaagtacaaggagatcttcttcgatcagtcgaagaacggctacgctgggtacattgacggcggggcctctcaggaggagttctacaagttcatcaagc
cgattctggagaagatggacggcacggaggagctgctggtgaagctcaatcgcgaggacctcctgaggaagcagcggacattcgataacggcagcatcccacacca
gattcatctcggggagctgcacgctatcctgaggaggcaggaggacttctaccctttcctcaaggataaccgcgagaagatcgagaagattctgactttcaggatcccg
tactacgtcggcccactcgctaggggcaactcccgcttcgcttggatgacccgcaagtcagaggagacgatcacgccgtggaacttcgaggaggtggtcgacaaggg
cgctagcgctcagtcgttcatcgagaggatgacgaatttcgacaagaacctgccaaatgagaaggtgctccctaagcactcgctcctgtacgagtacttcacagtctac
aacgagctgactaaggtgaagtatgtgaccgagggcatgaggaagccggctttcctgtctggggagcagaagaaggccatcgtggacctcctgttcaagaccaaccg
gaaggtcacggttaagcagctcaaggaggactacttcaagaagattgagtgcttcgattcggtcgagatctctggcgttgaggaccgcttcaacgcctccctggggacc
taccacgatctcctgaagatcattaaggataaggacttcctggacaacgaggagaatgaggatatcctcgaggacattgtgctgacactcactctgttcgaggaccgg
gagatgatcgaggagcgcctgaagacttacgcccatctcttcgatgacaaggtcatgaagcagctcaagaggaggaggtacaccggctggggggaggctgagcagga
agctcatcaacggcattcgggacaagcagtccgggaagacgatcctcgacttcctgaagagcgatggcttcgcgaaccgcaatttcatgcagctgattcacgatgaca
gcctcacattcaaggaggatatccagaaggctcaggtgagcggccaggggggactcgctgcacgagcatatcgcgaacctcgctggctcgccagctatcaagaaggg
gattctgcagaccgtgaaggttgtggacgagctggtgaaggtcatgggcaggcacaagcctgagaacatcgtcattgagatggcccgggagaatcagaccacgcag
aagggccagaagaactcacgcgagaggatgaagaggatcgaggagggcattaaggagctggggtcccagatcctcaaggagcacccggtggagaacacgcagct
gcagaatgagaagctctacctgtactacctccagaatggccgcgatatgtatgtggaccaggagctggatattaacaggctcagcgattacgacgtcgatgccatcgtt
ccacagtcattcctgaaggatgactccattgacaacaaggtcctcaccaggtcggacaagaaccggggcaagtctgataatgttccttcagaggaggtcgttaagaag
atgaagaactactggcgccagctcctgaatgccaagctgatcacgcagcggaagttcgataacctcacaaaggctgagaggggcgggctctctgagctggacaagg
cgggcttcatcaagaggcagctggtcgagacacggcagatcactaagcacgttgcgcagattctcgactcacggatgaacactaagtacgatgagaatgacaagctg
atccgcgaggtgaaggtcatcaccctgaagtcaaagctcgtctccgacttcaggaaggatttccagttctacaaggttcgggagatcaacaattaccaccatgcccatg
acgcgtacctgaacgcggtggtcggcacagctctgatcaagaagtacccaaagctcgagagcgagttcgtgtacggggactacaaggtttacgatgtgaggaagatg
atcgccaagtcggagcaggagattggcaaggctaccgccaagtacttcttctactctaacattatgaatttcttcaagacagagatcactctggccaatggcgagatcc
ggaagcgccccctcatcgagacgaacggcgagacgggggagatcgtgtgggacaagggcagggatttcgcgaccgtcaggaaggttctctccatgccacaagtgaa
tatcgtcaagaagacagaggtccagactggcgggttctctaaggagtcaattctgcctaagcggaacagcgacaagctcatcgcccgcaagaaggactgggatccga
agaagtacggcgggttcgacagccccactgtggcctactcggtcctggttgtggcgaaggttgagaagggcaagtccaagaagctcaagagcgtgaaggagctgct
ggggatcacgattatggagcgctccagcttcgagaagaacccgatcgatttcctggaggcgaagggctacaaggaggtgaagaaggacctgatcattaagctcccca
agtactcactcttcgagctggagaacggcaggaagcggatgctggcttccgctggcgagctgcagaaggggaacgagctggctctgccgtccaagtatgtgaacttcc
tctacctggcctcccactacgagaagctcaagggcagccccgaggacaacgagcagaagcagctgttcgtcgagcagcacaagcattacctcgacgagatcattgag
cagatttccgagttctccaagcgcgtgatcctggccgacgcgaatctggataaggtcctctccgcgtacaacaagcaccgcgacaagccaatcagggagcaggctga
gaatatcattcatctcttcaccctgacgaacctcggcgcccctgctgctttcaagtacttcgacacaactatcgatcgcaagaggtacacaagcactaaggaggtcctg
gacgcgaccctcatccaccagtcgattaccggcctctacgagacgcgcatcgacctgtctcagctcgggggcgactcaggcggctcatcgggcgggtcaagcgggtc
ggagacaccgggcacatcagagagcgctacccctgagtcatcaggcggctcttcaggcggcagctcaaccctgaacattgaggacgagtaccggctgcacgagacg
agcaaggagccagacgtttcgctcggcagcacttggctctctgacttcccacaggcttgggccgagactggcggcatgggcctggccgtgcgccaggctccactgatc
atccctctgaaggcgacctccaccccggtttctattaagcagtacccgatgagccaggaggccaggctggggatcaagccacacattcagcggctgctggaccaggg
catcctggtgccatgccagtccccgtggaatactccgctcctgccggtgaagaagcctgggacaaacgactacaggccggttcaggatctcagggaggtgaacaagc
gcgtggaggacatccatccgacagtgccgaacccgtacaatctgctgtcgggcctgcctccgagccaccagtggtacaccgtcctggacctcaaggacgctttcttctg
cctgcggctgcacccgacgtctcagccgctgttcgcgttcgagtggcgcgacccagagatgggcatttccggccagctgacctggacacgcctaccccagggcttcaa
gaactccccgactctcttcaacgaggctctccaccgggatctcgcggacttcaggattcagcatcccgatctgatcctgctccagtatgttgacgacctcctcctggccgc
gacgtcggagctggactgccagcagggcacccgggcgctgctgcagacactgggcaatctggggtaccgcgcctctgcgaagaaggcgcagatctgccagaagca
agtgaagtacctgggctacctcctgaaggagggccagcgctggctcactgaggcgaggaaggagactgttatggccagcccactccaaagactccgaggcagctc
agggagttcctcggcaaggctgggttctgccgcctgttcatccctgggttcgctgagatggctgcgccgctctacccgctgactaagccggggacactgttcaactggg

ggccagaccagcagaaggcgtaccaggagattaagcaggcgctgctgacggccccagcgctcggcctaccagacctgacgaagccgttcgagctgttcgttgacga
gaagcaggggtacgcgaagggcgtgctgacacagaagctggggccttggcgccgcccggtcgcgtacctgtcgaagaagctggacccagtcgctgctgggtggcct
ccatgcctccggatggtcgctgctattgcggttctgaccaaggatgcggggaagctcacaatggggcagcctctcgtgatcctggctccacatgcggtggaggcgctg
gtgaagcagccaccggaccggtggctgtcgaacgctcggatgacacactaccaggcgctcctcctcgatacagaccgggttcagttcgggcctgtggttgctctgaac
ccagccacactgctgccactccctgaggagggcctccagcacaattgcctcgacatcctggctgaggcgcacggcacccgccctgatctcaccgaccagcctctgcca
gatgctgaccacacctggtacacggatgggtcctcgctgctgcaggagggccagaggaaggcgggcgccgccgtcaccacagagacagaggttatttgggccaagg
ccctaccggctggcaccagcgcccagcgcgctgagctgatcgcgctgactcaggcgctgaagatggccgaggggaagaagctcaatgtttacaccgactcgcggtac
gcgttcgctacagctcacattcatggggagatctaccgccggcgcgggtggctgacttcggagggcaaggagattaagaataaggacgagatcctggccctgctcaa
ggcgctgttcctgccgaagcgcctctcaatcattcactgcccgggccaccagaagggccattcggccgaggctaggggcaatcggatggctgaccaggcggcgcgga
aggcggctatcaccgagactcccgatacatctaccctcctgatcgagaactcgagcccaagcggcgggagcaagcggactgcggatgggtctgagttcgagccaaa
gaagaagaggaaggtgtga

**Synthetic P165S-1 for generation of pZ1PE3b**

BsaI-T1P-sgR-rtT-PBS-OsU3t-TaU3p-T2P-BsaI

GGTCTCTGGCGCGGTGCCAATCATGCGTCGGTTTTAGAGCTAGAAATAGCAAGTTAAAATAAGGCTAGTCCGTTATCAACTTGAAAAAGTGGCACC
GAGTCGGTGCGGACAGGTGAGTCGACGCATGATTGTTTTTTTTTTCGTTTTGCATTGAGTTTTCTCCGTCGCATGTTTGCAGCATGAATCCAAACCACA
CGGAGTTCAAATTCCCACAGATTAAGGCTCGTCCGTCGCACAAGGTAATGTGTGAATATTATATCTGTCGTGCAAAATTGCCTGGCCTGCACAATTGCTG
TTATAGTTGGCGGCAGGGAGAGTTTTAACATTGACTAGCGTGCTGATAATTTGTGAGAAATAATAATTGACAAGTAGATACTGACATTTGAGAAGAGCT
TCTGAACTGTTATTAGTAACAAAAATGGAAAGCTGATGCACGGAAAAAGGAAAGAAAAAGCCATACTTTTTTTTAGGTAGGAAAAGAAAAAGCCATAC
GAGACTGATGTCTCTCAGATGGGCCGGGATCTGTCTATCTAGCAGGCAGCAGCCCACCAACCTCACGGGCCAGCAATTACGAGTCCTTCTAAAAGCTC
CCGCCGAGGGGCGCTGGCGCTGCTGTGCAGCAGCACGTCTAACATTAGTCCCACCTCGCCAGTTTACAGGGAGCAGAACCAGCTTATAAGCGGAGGC
GCGGCACCAAGAAGCGAGGTGAGTCGACGCATGATGTTTAGAGACC

**Synthetic P165S-2 for generation of pZ1PE3**

BsaI-T1P-sgR-rtT-PBS-OsU3t-TaU3p-T2P-BsaI

GGTCTCTGGCGCGGTGCCAATCATGCGTCGGTTTTAGAGCTAGAAATAGCAAGTTAAAATAAGGCTAGTCCGTTATCAACTTGAAAAAGTGGCACC
GAGTCGGTGCGGACAGGTGagtCGACGCATGATTGGCATTTTTTTTTTCGTTTTGCATTGAGTTTTCTCCGTCGCATGTTTGCAGCATGAATCCAAACC
ACACGGAGTTCAAATTCCCACAGATTAAGGCTCGTCCGTCGCACAAGGTAATGTGTGAATATTATATCTGTCGTGCAAAATTGCCTGGCCTGCACAATTG
CTGTTATAGTTGGCGGCAGGGAGAGTTTTAACATTGACTAGCGTGCTGATAATTTGTGAGAAATAATAATTGACAAGTAGATACTGACATTTGAGAAGA
GCTTCTGAACTGTTATTAGTAACAAAAATGGAAAGCTGATGCACGGAAAAAGGAAAGAAAAAGCCATACTTTTTTTTAGGTAGGAAAAGAAAAAGCC
ATACGAGACTGATGTCTCTCAGATGGGCCGGGATCTGTCTATCTAGCAGGCAGCAGCCCACCAACCTCACGGGCCAGCAATTACGAGTCCTTCTAAAA
GCTCCCGCCGAGGGGCGCTGGCGCTGCTGTGCAGCAGCACGTCTAACATTAGTCCCACCTCGCCAGTTTACAGGGAGCAGAACCAGCTTATAAGCGG
AGGCGCGGCACCAAGAAGCGCAGGAGACGCCCATCGTCGGTTTAGAGACC

**Synthetic W542L for generation of pG3R2R3-W542L (pZ1WS/pZ1WS-Csy4)**

BsaI-T1W-sgR-rtT-PBS-OsU3t-TaU3p-T2W-BsaI

GGTCTCTGGCGGGGATGGTGGTGCAGTGGGGTTTTAGAGCTAGAAATAGCAAGTTAAAATAAGGCTAGTCCGTTATCAACTTGAAAAAGTGGCAC
CGAGTCGGTGCTAGAACCTGTCTTCTAACTGCACCACCATTTTTTTTTTTCGTTTTGCATTGAGTTTTCTCCGTCGCATGTTTGCAGCATGAATCCAAAC
CACACGGAGTTCAAATTCCCACAGATTAAGGCTCGTCCGTCGCACAAGGTAATGTGTGAATATTATATCTGTCGTGCAAAATTGCCTGGCCTGCACAATT
GCTGTTATAGTTGGCGGCAGGGAGAGTTTTAACATTGACTAGCGTGCTGATAATTTGTGAGAAATAATAATTGACAAGTAGATACTGACATTTGAGAAG
AGCTTCTGAACTGTTATTAGTAACAAAAATGGAAAGCTGATGCACGGAAAAAGGAAAGAAAAAGCCATACTTTTTTTTAGGTAGGAAAAGAAAAAGC
CATACGAGACTGATGTCTCTCAGATGGGCCGGGATCTGTCTATCTAGCAGGCAGCAGCCCACCAACCTCACGGGCCAGCAATTACGAGTCCTTCTAAAA
GCTCCCGCCGAGGGGCGCTGGCGCTGCTGTGCAGCAGCACGTCTAACATTAGTCCCACCTCGCCAGTTTACAGGGAGCAGAACCAGCTTATAAGCGG
AGGCGCGGCACCAAGAAGCGTAACTGCACCACCATCCCGGTTTAGAGACC

**Synthetic WS-Csy4 for generation of pL2L1-WS-Csy4 (pZ1WS-Csy4)**

BsaI-T1W-sgR-rtT-PBS-HDV-Csy4-T1S-sgR-rtT-PBS-BsaI

GGTCTCATGCACGGGATGGTGGTGCAGTGGGGTTTTAGAGCTAGAAATAGCAAGTTAAAATAAGGCTAGTCCGTTATCAACTTGAAAAAGTGGCA
CCGAGTCGGTGCTAGAACCTGTCTTCTAACTGCACCACCATGGCCGGCATGGTCCCAGCCTCCTCGCTGGCGCCGGCTGGGCAACATGCTTCGGCAT
GGCGAATGGGACGTTCACTGCCGTATAGGCAGCCTTGAAAGCCCCACCACTAGTTTTAGAGCTAGAAATAGCAAGTTAAAATAAGGCTAGTCCGTT
ATCAACTTGAAAAAGTGGCACCGAGTCGGTGCTGCCTATGATACCTATTGGTGGGGCTTTCGGCCAGAGACC

##### Synthetic WS-pegR for generation of pL2L1-WS-pegR (pZ1WS)

BsaI-T1W-sgR-rtT-PBS-HDV-tMet-T1S-sgR-rtT-PBS-BsaI

GGTCTCATGCACGGGATGGTGGTGCAGTGGGGTTTTAGAGCTAGAAATAGCAAGTTAAAATAAGGCTAGTCCGTTATCAACTTGAAAAAGTGGCA
CCGAGTCGGTGCTAGAACCTGTCTTCTAACTGCACCACCATGGCCGGCATGGTCCCAGCCTCCTCGCTGGCGCCGGCTGGGCAACATGCTTCGGCAT
GGCGAATGGGACAACAACAAATCAGAGTGGCGCAGCGGAAGCGTGGTGGGCCCATAACCCACAGGTCCCAGGATCGAAACCTGGCTCTGATACCT
TGAAAGCCCCACCACTAGTTTTAGAGCTAGAAATAGCAAGTTAAAATAAGGCTAGTCCGTTATCAACTTGAAAAAGTGGCACCGAGTCGGTGCTGC
CTATGATACCTATTGGTGGGGCTTTCGGCCAGAGACC

##### Synthetic WS-sgR for generation of pR1R4-WS-sgR (pZ1WS/pZ1WS-Csy4)

BsaI-T2W-sgR-OsU3t-TaU3p-T2S-BsaI

GGTCTCTGGCGTAACTGCACCACCATCCCGGTTTTAGAGCTAGAAATAGCAAGTTAAAATAAGGCTAGTCCGTTATCAACTTGAAAAAGTGGCACCG
AGTCGGTGCTTTTTTTTTTCGTTTTGCATTGAGTTTTCTCCGTCGCATGTTTGCAGCATGAATCCAAACCACACGGAGTTCAAATTCCCACAGATTAAGG
CTCGTCCGTCGCACAAGGTAATGTGTGAATATTATATCTGTCGTGCAAAATTGCCTGGCCTGCACAATTGCTGTTATAGTTGGCGGCAGGGAGAGTTTTA
ACATTGACTAGCGTGCTGATAATTTGTGAGAAATAATAATTGACAAGTAGATACTGACATTTGAGAAGAGCTTCTGAACTGTTATTAGTAACAAAAATGG
AAAGCTGATGCACGGAAAAAGGAAAGAAAAAGCCATACTTTTTTTTAGGTAGGAAAAGAAAAAGCCATACGAGACTGATGTCTCTCAGATGGGCCGG
GATCTGTCTATCTAGCAGGCAGCAGCCCACCAACCTCACGGGCCAGCAATTACGAGTCCTTCTAAAAGCTCCCGCCGAGGGGCGCTGGCGCTGCTGTG
CAGCAGCACGTCTAACATTAGTCCCACCTCGCCAGTTTACAGGGAGCAGAACCAGCTTATAAGCGGAGGCGCGGCACCAAGAAGCGATATGATCCTG
GATGGTGAGTTTAGAGACC

##### Synthetic S621I for generation of pL4L3-S621I (pZ1WS/pZ1WS-Csy4)

BsaI-T1S-sgR-rtT-PBS-OsU3t-TaU3p-T2S-BsaI

GGTCTCTGGCGCTTGAAAGCCCCACCACTAGTTTTAGAGCTAGAAATAGCAAGTTAAAATAAGGCTAGTCCGTTATCAACTTGAAAAAGTGGCACC
GAGTCGGTGCTGCCTATGATACCTATTGGTGGGGCTTTCTTTTTTTTTTCGTTTTGCATTGAGTTTTCTCCGTCGCATGTTTGCAGCATGAATCCAAACC
ACACGGAGTTCAAATTCCCACAGATTAAGGCTCGTCCGTCGCACAAGGTAATGTGTGAATATTATATCTGTCGTGCAAAATTGCCTGGCCTGCACAATTG
CTGTTATAGTTGGCGGCAGGGAGAGTTTTAACATTGACTAGCGTGCTGATAATTTGTGAGAAATAATAATTGACAAGTAGATACTGACATTTGAGAAGA
GCTTCTGAACTGTTATTAGTAACAAAAATGGAAAGCTGATGCACGGAAAAAGGAAAGAAAAAGCCATACTTTTTTTTAGGTAGGAAAAGAAAAAGCC
ATACGAGACTGATGTCTCTCAGATGGGCCGGGATCTGTCTATCTAGCAGGCAGCAGCCCACCAACCTCACGGGCCAGCAATTACGAGTCCTTCTAAAA
GCTCCCGCCGAGGGGCGCTGGCGCTGCTGTGCAGCAGCACGTCTAACATTAGTCCCACCTCGCCAGTTTACAGGGAGCAGAACCAGCTTATAAGCGG
AGGCGCGGCACCAAGAAGCGATATGATCCTGGATGGTGAGTTTAGAGACC

##### Synthetic OsALS-1pegR for generation of p35C-ALS-S1 and pU3-ALS-S1

BsaI-Spacer-sgR-rtT-PBS-BsaI

GGTCTCATGCAGTGCTGCCTATGATCCCAAGGTTTTAGAGCTAGAAATAGCAAGTTAAAATAAGGCTAGTCCGTTATCAACTTGAAAAAGTGGCACC
GAGTCGGTGCTGAATGCGCCCCCAaTTGGGATCATAGGGCCAGAGACC

##### Synthetic OsGAPDH-1pegR for generation of p35C-GAPDH and pU3-GAPDH

BsaI-Spacer-sgR-rtT-PBS-BsaI

GGTCTCATGCAGAGTATGTCGTGGAGTCCACGTTTTAGAGCTAGAAATAGCAAGTTAAAATAAGGCTAGTCCGTTATCAACTTGAAAAAGTGGCAC
CGAGTCGGTGCAGTGAAGACACCGtTGGACTCCACGACAGGCCAGAGACC

##### Synthetic OsALS-2pegR1 for generation of p35C-ALS-WS and pL2L1-ALS-WS

BsaI-Spacer-sgR-rtT-PBS-HDV-tMet-Spacer-sgR-rtT-PBS-BsaI

GGTCTCATGCAGGGTATGGTGGTGCAATGGGGTTTTAGAGCTAGAAATAGCAAGTTAAAATAAGGCTAGTCCGTTATCAACTTGAAAAAGTGGCAC
CGAGTCGGTGCAAACCTATCTTCTAATTGCACCACCATGGCCGGCATGGTCCCAGCCTCCTCGCTGGCGCCGGCTGGGCAACATGCTTCGGCATGG
CGAATGGGACAACAACAAATCAGAGTGGCGCAGCGGAAGCGTGGTGGGCCCATAACCCACAGGTCCCAGGATCGAAACCTGGCTCTGATACCTTG
AATGCGCCCCCACTTGTTTTAGAGCTAGAAATAGCAAGTTAAAATAAGGCTAGTCCGTTATCAACTTGAAAAAGTGGCACCGAGTCGGTGCTGCCTA
TGATACCAATTGGGGGCGCATTCGGCCAGAGACC

**Synthetic OsALS-2pegR2 for generation of p2xU3-ALS-WS, pG3R23-OsWS, pR1R4-OsWS, and pL4L3-OsWS**

BsaI-Spacer-sgR-rtT-PBS-HDV-OsU3t-TaU3p-tMet-Spacer-sgR-rtT-PBS-BsaI

GGTCTCATGCAGGGTATGGTGGTGCAATGGGGTTTTAGAGCTAGAAATAGCAAGTTAAAATAAGGCTAGTCCGTTATCAACTTGAAAAAGTGGCAC
CGAGTCGGTGCAAACCTATCTTCTAATTGCACCACCATGGCCGGCATGGTCCCAGCCTCCTCGCTGGCGCCGGCTGGGCAACATGCTTCGGCATGG
CGAATGGGACTTTTTTTTTTCGTTTTGCATTGAGTTTTCTCCGTCGCATGTTTGCAGTTTTATTTTCCGTTTTGCATTGAAATTTCTCCGTCTCATGTTTG
C**AG**CGTGTTCAACATGAATCCAAACCACACGGAGTTCAAATTCCCACAGATTAAGGCTCGTCCGTCGCACAAGGTAATGTGTGAATATTATATCTGT
CGTGCAAAATTGCCTGGCCTGCACAATTGCTGTTATAGTTGGCGGCAGGGAGAGTTTTAACATTGACTAGCGTGCTGATAATTTGTGAGAAATAAT
AATTGACAAGTAGATACTGACATTTGAGAAGAGCTTCTGAACTGTTATTAGTAACAAAAATGGAAAGCTGATGCACGGAAAAAGGAAAGAAAAAG
CCATACTTTTTTTTAGGTAGGAAAAGAAAAAGCCATACGAGACTGATGTCTCTCAGATGGGCCGGGATCTGTCTATCTAGCAGGCAGCAGCCCACC
AACCTCACGGGCCAGCAATTACGAGTCCTTCTAAAAGCTCCCGCCGAGGGGCGCTGGCGCTGCTGTGCAGCAGCACGTCTAACATTAGTCCCACCT
CGCCAGTTTACAGGGAGCAGAACCAGCTTATAAGCGGAGGCGCGGCACCAAGAAGCGAACAACAAAACAACAAATCAGAGTGGCGCAGCGGAAG
CGTGGTGGGCCCATAACCCACAGGTCCCAGGATCGAAACCTGGCTCTGATACCTTGAATGCGCCCCCACTTGTTTTAGAGCTAGAAATAGCAAGTTA
AAATAAGGCTAGTCCGTTATCAACTTGAAAAAGTGGCACCGAGTCGGTGCTGCCTATGATACCAATTGGGGGCGCATTCGGCCAGAGACC

**The pegRNA and sgRNA cassettes in pZ1PE3, pZ1PE3b, and pZ1WS et al.**

OsU3p-T1-sgR-rtT-PBS-OsU3t-TaU3p-T2-sgR-TaU3t

AGTAATTCATCCAGGTCACCAAGTTCTAGGATTTTCAGAACTGCAACTTATTTTATCAAGGAATCTTTAAACATACGAACAGATCACTTAAAGTTCTTCTG
AAGCAACTTAAAGTTATCAGGCTTGCATGGATCTTGGAGGAATCAGATGTGCAGTCAGGGACCATAGCACAAGACAGGCGTCTTCTACTGGTGCTACC
AGCAAATGCTGGAAGCCGGGAACACTGGGTACGTTGGAAACCACGTGATGTGAAGAAGTAAGATAAACTGTAGGAGAAAAGCATTTCGTAGTGGGC
CATGAAGCCTTTCAGGACATGTATTGCAGTATGGGCCGGCCCATTACGCAATTGGACGACAACAAAGTCTAGTATTAGTACCACCTCGGCTATCCACATA
GATCAAAGCTGATTTAAAAGAGTTGTGCAGATGATCCGTGGCGNNNNNNNNNNNNNNNNNNNGTTTTAGAGCTAGAAATAGCAAGTTAAAATAA
GGCTAGTCCGTTATCAACTTGAAAAAGTGGCACCGAGTCGGTGCNNNNNNNNNNNNNNNNNNNNNNNNNNNNNTTTTTTTTTTCGTTTTGCAT
TGAGTTTTCTCCGTCGCATGTTTGCAGCATGAATCCAAACCACACGGAGTTCAAATTCCCACAGATTAAGGCTCGTCCGTCGCACAAGGTAATGTGTGA
ATATTATATCTGTCGTGCAAAATTGCCTGGCCTGCACAATTGCTGTTATAGTTGGCGGCAGGGAGAGTTTTAACATTGACTAGCGTGCTGATAATTTGTG
AGAAATAATAATTGACAAGTAGATACTGACATTTGAGAAGAGCTTCTGAACTGTTATTAGTAACAAAAATGGAAAGCTGATGCACGGAAAAAGGAAAG
AAAAAGCCATACTTTTTTTTAGGTAGGAAAAGAAAAAGCCATACGAGACTGATGTCTCTCAGATGGGCCGGGATCTGTCTATCTAGCAGGCAGCAGCC
CACCAACCTCACGGGCCAGCAATTACGAGTCCTTCTAAAAGCTCCCGCCGAGGGGCGCTGGCGCTGCTGTGCAGCAGCACGTCTAACATTAGTCCCAC
CTCGCCAGTTTACAGGGAGCAGAACCAGCTTATAAGCGGAGGCGCGGCACCAAGAAGCGNNNNNNNNNNNNNNNNNNNGTTTTAGAGCTAGAA
ATAGCAAGTTAAAATAAGGCTAGTCCGTTATCAACTTGAAAAAGTGGCACCGAGTCGGTGCTTTTTTTTTTGTCCTTCTGTTTTTTTAGTCAGTCTCTTTT
TTCAGAAGTACAACATCTT

Note: the underlined part comes from the synthetic fragments and cassettes with the same structure as this one are harbored in pZ1PE3, pZ1PE3b, pG3R2R3-W542L, pL4L3-S621I, pZ1WS, and pZ1WS-Csy4.

**The two pegRNA cassettes in pL2L1-WS-Csy4 and pZ1WS-Csy4**

35S-CmYLCV-Csy4-P2A-ZsGreen-polyA-Csy4-T1W-sgR-rtT-PBS-HDV-Csy4-T1S-sgR-rtT-PBS-HDV-Csy4-HSPt

ATGGAGTCAAAGATTCAAATAGAGGACCTAACAGAACTCGCCGTAAAGACTGGCGAACAGTTCATACAGAGTCTCTTACGACTCAATGACAAGAAGAA
AATCTTCGTCAACATGGTGGAGCACGACACACTTGTCTACTCCAAAAATATCAAAGATACAGTCTCAGAAGACCAAAGGGCAATTGAGACTTTTCAACA
AAGGGTAATATCCGGAAACCTCCTCGGATTCCATTGCCCAGCTATCTGTCACTTTATTGTGAAGATAGTGGAAAAGGAAGGTGGCTCCTACAAATGCCA
TCATTGCGATAAAGGAAAGGCCATCGTTGAAGATGCCTCTGCCGACAGTGGTCCCAAAGATGGACCCCCACCCACGAGGAGCATCGTGGAAAAAGAA
GACGTTCCAACCACGTCTTCAAAGCAAGTGGATTGATGTGATTGGCAGACATACTGTCCCACAAATGAAGATGGAATCTGTAAAAGAAAACGCGTGAA
ATAATGCGTCTGACAAAGGTTAGGTCGGCTGCCTTTAATCAATACCAAAGTGGTCCCTACCACGATGGAAAAACTGTGCAGTCGGTTTGGCTTTTTCTG
ACGAACAAATAAGATTCGTGGCCGACAGGTGGGGGTCCACCATGTGAAGGCATCTTCAGACTCCAATAATGGAGCAATGACGTAAGGGCTTACGAAA
TAAGTAAGGGTAGTTTGGGAAATGTCCACTCACCCGTCAGTCTATAAATACTTAGCCCCTCCCTCATTGTTAAGGGAGCAAAATCTCAGAGAGATAGTCC
TAGAGAGAGAAAGAGAGCAAGTAGCCTAGAAGTAGTCAAGGCGGCGAAGTATTCAGGCACGTGGCCAGGAAGAAGAAAAGCCAAGACGACGAAA
ACAGGTAAGAGCTAAGCATCTAGATGGACCACTACCTGGACATCCGGCTCCGCCCGGACCCCGAGTTCCCGCCCGCCCAGCTGATGTCCGTGCTCTTC
GGCAAGCTGCACCAGGCCCTCGTGGCCCAGGGCGGCGACCGCATCGGCGTCTCCTTCCCGGACCTGGACGAGTCCAGGAGCCGCCTGGGCGAGAG
GCTCCGCATCCACGCCAGCGCGGACGACCTGAGGGCCCTCCTCGCCCGCCCGTGGCTGGAGGGCCTCAGGGACCACCTCCAGTTCGGCGAGCCGGC
CGTGGTCCCCCACCCAACCCCGTACCGCCAGGTGTCCCGCGTGCAGGCCAAGTCCAACCCCGAGAGGCTGCGCAGGCGCCTCATGAGGCGCCACGA
CCTGTCGGAGGAGGAGGCTAGGAAGCGCATCCCCGACACCGTGGCTAGGGCGCTGGACCTCCCCTTCGTCACCCTCCGCTCCCAGAGCACGGGCCA
GCACTTCCGGCTGTTCATCCGCCACGGCCCACTCCAGGTGACCGCCGAGGAGGGCGGCTTCACGTGCTACGGCCTCTCCAAGGGCGGCTTCGTCCCG
TGGTTCGGGCTCCGGCGCCACCAACTTCTCGCTGCTGAAGCAGGCGGGCGACGTGGAGGAGAACCCGGGCCCGATGGCCCAGTCCAAGCACGGCC
TGACCAAGGAGATGACGATGAAGTACCGCATGGAGGGCTGCGTGGACGGCCACAAGTTCGTCATCACCGGCGAGGGCATCGGCTACCCGTTCAAGG
GCAAGCAGGCCATCAACCTCTGCGTGGTGGAGGGCGGCCCCCTGCCGTTCGCCGAGGACATCCTCAGCGCCGCGTTCATGTACGGCAACCGCGTGTT
CACCGAGTACCCGCAGGACATCGTGGACTACTTCAAGAACTCCTGCCCAGCCGGCTACACGTGGGACAGGAGCTTCCTGTTCGAGGACGGCGCCGTG
TGCATCTGCAACGCGGACATCACCGTGTCCGTCGAGGAGAACTGCATGTACCACGAGAGCAAGTTCTACGGCGTGAACTTCCCGGCGGACGGCCCCG
TCATGAAGAAGATGACGGACAACTGGGAGCCGTCCTGCGAGAAGATCATCCCGGTGCCCAAGCAGGGCATCCTGAAGGGCGACGTCAGCATGTACCT

CCTGCTCAAGGACGGCGGCAGGCTCAGGTGCCAGTTCGACACGGTGTACAAGGCGAAGTCCGTCCCGAGGAAGATGCCCGACTGGCACTTCATCCA
GCACAAGCTCACCCGCGAGGACAGGAGCGACGCCAAGAACCAGAAGTGGCACCTGACCGAGCACGCTATCGCCTCCGGCAGCGCGCTCCCCTGAGC
TCAAAAAAAAAAAAAAAAAAAAAAAAAAAAAAAAAAAAAAAAAAAAAAAAAAAAGAATTGGTACC**GTTCACTGCCGTATAGGCAG**CGGGATGGTGGT
GCAGTGGGGTTTTAGAGCTAGAAATAGCAAGTTAAAATAAGGCTAGTCCGTTATCAACTTGAAAAAGTGGCACCGAGTCGGTGCTAGAACCTGTCT
TCTAACTGCACCACCATGGCCGGCATGGTCCCAGCCTCCTCGCTGGCGCCGGCTGGGCAACATGCTTCGGCATGGCGAATGGGACGTTCACTGCCG
TATAGGCAGCCTTGAAAGCCCCACCACTAGTTTTAGAGCTAGAAATAGCAAGTTAAAATAAGGCTAGTCCGTTATCAACTTGAAAAAGTGGCACCG
AGTCGGTGCTGCCTATGATACCTATTGGTGGGGCTTTCGGCCGGCATGGTCCCAGCCTCCTCGCTGGCGCCGGCTGGGCAACATGCTTCGGCATGGC
GAATGGGACGTTCACTGCCGTATAGGCAGCCTAGGGATATCTCCGGGGCTAATTGAATATGAAGATGAAGATGAAATATTTGGTGTGTCAAATAAAAAG
CTGGTGTGCTTAAGTTTGTGTTTTTTTCTTGGCTTGTTGTGTTATGAATTTGTGGCTTTTTCTAATATTAAATGAATGTAAGATCTCATTATAATGAATAAAC
AAATGTTTCTATAATCCATTGTGAATGTTTTGTTGGATCTCTTCTGCAGCATATAACTACTGTATGTGCTATGGTATGGACTATGGAATATGATTAAAGATAA
G

Note: the underlined part comes from the synthetic fragment.

##### The two pegRNA cassettes in pL2L1-WS-pegR and pZ1WS

35S-CmYLCV-U6-tGly-T1W-sgR-rtT-PBS-HDV-tMet-T1S-sgR-rtT-PBS-HDV-polyT-HSPt

ATGGAGTCAAAGATTCAAATAGAGGACCTAACAGAACTCGCCGTAAAGACTGGCGAACAGTTCATACAGAGTCTCTTACGACTCAATGACAAGAAGAA
AATCTTCGTCAACATGGTGGAGCACGACACACTTGTCTACTCCAAAAATATCAAAGATACAGTCTCAGAAGACCAAAGGGCAATTGAGACTTTTCAACA
AAGGGTAATATCCGGAAACCTCCTCGGATTCCATTGCCCAGCTATCTGTCACTTTATTGTGAAGATAGTGGAAAAGGAAGGTGGCTCCTACAAATGCCA
TCATTGCGATAAAGGAAAGGCCATCGTTGAAGATGCCTCTGCCGACAGTGGTCCCAAAGATGGACCCCCACCCACGAGGAGCATCGTGGAAAAAGAA
GACGTTCCAACCACGTCTTCAAAGCAAGTGGATTGATGTGATTGGCAGACATACTGTCCCACAAATGAAGATGGAATCTGTAAAAGAAAACGCGTGAA
ATAATGCGTCTGACAAAGGTTAGGTCGGCTGCCTTTAATCAATACCAAAGTGGTCCCTACCACGATGGAAAAACTGTGCAGTCGGTTTGGCTTTTTCTG
ACGAACAAATAAGATTCGTGGCCGACAGGTGGGGGTCCACCATGTGAAGGCATCTTCAGACTCCAATAATGGAGCAATGACGTAAGGGCTTACGAAA
TAAGTAAGGGTAGTTTGGGAAATGTCCACTCACCCGTCAGTCTATAAATACTTAGCCCCTCCCTCATTGTTAAGGGAGCAAAATCTCAGAGAGATAGTCC
TAGAGAGAGAAAGAGAGCAAGTAGCCTAGAAGTAGTCAAGGCGGCGAAGTATTCAGGCACGTGGCCAGGAAGAAGAAAAGCCAAGACGACGAAA
ACAGGTAAGAGCTAAGCATCTAGAAAGTTGAAAACAATCTTCAAAAGTCCCACATCGCTTAGATAAGAAAACGAAGCTGAGTTTATATACAGCTAGAGT
CGAAGTAGTGATTGAACAAAGCACCAGTGGTCTAGTGGTAGAATAGTACCCTGCCACGGTACAGACCCGGGTTCGATTCCCGGCTGGTGCACGGGAT
GGTGGTGCAGTGGGGTTTTAGAGCTAGAAATAGCAAGTTAAAATAAGGCTAGTCCGTTATCAACTTGAAAAAGTGGCACCGAGTCGGTGCTAGAA
CCTGTCTTCTAACTGCACCACCATGGCCGGCATGGTCCCAGCCTCCTCGCTGGCGCCGGCTGGGCAACATGCTTCGGCATGGCGAATGGGACAACA
ACAAATCAGAGTGGCGCAGCGGAAGCGTGGTGGGCCCATAACCCACAGGTCCCAGGATCGAAACCTGGCTCTGATACCTTGAAAGCCCCACCACT
AGTTTTAGAGCTAGAAATAGCAAGTTAAAATAAGGCTAGTCCGTTATCAACTTGAAAAAGTGGCACCGAGTCGGTGCTGCCTATGATACCTATTGGT
GGGGCTTTCGGCCGGCATGGTCCCAGCCTCCTCGCTGGCGCCGGCTGGGCAACATGCTTCGGCATGGCGAATGGGACTTTTTTTTGATATCTCCGGGG
CTAATTGAATATGAAGATGAAGATGAAATATTTGGTGTGTCAAATAAAAAGCTGGTGTGCTTAAGTTTGTGTTTTTTTCTTGGCTTGTTGTGTTATGAATT
TGTGGCTTTTTCTAATATTAAATGAATGTAAGATCTCATTATAATGAATAAACAAATGTTTCTATAATCCATTGTGAATGTTTTGTTGGATCTCTTCTGCAGC
ATATAACTACTGTATGTGCTATGGTATGGACTATGGAATATGATTAAAGATAAG

Note: the underlined part comes from the synthetic fragment.

##### The two sgRNA cassettes in pR1R4-WS-sgR, pZ1WS, and pZ1WS-Csy4

OsU3p-T2W-sgR-OsU3t-TaU3p-T2S-sgR-TaU3t

AGTAATTCATCCAGGTCACCAAGTTCTAGGATTTTCAGAACTGCAACTTATTTTATCAAGGAATCTTTAAACATACGAACAGATCACTTAAAGTTCTTCTG
AAGCAACTTAAAGTTATCAGGCTTGCATGGATCTTGGAGGAATCAGATGTGCAGTCAGGGACCATAGCACAAGACAGGCGTCTTCTACTGGTGCTACC
AGCAAATGCTGGAAGCCGGGAACACTGGGTACGTTGGAAACCACGTGATGTGAAGAAGTAAGATAAACTGTAGGAGAAAAGCATTTCGTAGTGGGC
CATGAAGCCTTTCAGGACATGTATTGCAGTATGGGCCGGCCCATTACGCAATTGGACGACAACAAAGTCTAGTATTAGTACCACCTCGGCTATCCACATA
GATCAAAGCTGATTTAAAAGAGTTGTGCAGATGATCCGTGGCGTAACTGCACCACCATCCCGGTTTTAGAGCTAGAAATAGCAAGTTAAAATAAGGC
TAGTCCGTTATCAACTTGAAAAAGTGGCACCGAGTCGGTGCTTTTTTTTTTCGTTTTGCATTGAGTTTTCTCCGTCGCATGTTTGCAGCATGAATCCAA
ACCACACGGAGTTCAAATTCCCACAGATTAAGGCTCGTCCGTCGCACAAGGTAATGTGTGAATATTATATCTGTCGTGCAAAATTGCCTGGCCTGCACAA
TTGCTGTTATAGTTGGCGGCAGGGAGAGTTTTAACATTGACTAGCGTGCTGATAATTTGTGAGAAATAATAATTGACAAGTAGATACTGACATTTGAGA
AGAGCTTCTGAACTGTTATTAGTAACAAAAATGGAAAGCTGATGCACGGAAAAAGGAAAGAAAAAGCCATACTTTTTTTTAGGTAGGAAAAGAAAAA
GCCATACGAGACTGATGTCTCTCAGATGGGCCGGGATCTGTCTATCTAGCAGGCAGCAGCCCACCAACCTCACGGGCCAGCAATTACGAGTCCTTCTAA
AAGCTCCCGCCGAGGGGCGCTGGCGCTGCTGTGCAGCAGCACGTCTAACATTAGTCCCACCTCGCCAGTTTACAGGGAGCAGAACCAGCTTATAAGC
GGAGGCGCGGCACCAAGAAGCGATATGATCCTGGATGGTGAGTTTTAGAGCTAGAAATAGCAAGTTAAAATAAGGCTAGTCCGTTATCAACTTGAAA
AAGTGGCACCGAGTCGGTGCTTTTTTTTTTGTCCTTCTGTTTTTTTAGTCAGTCTCTTTTTTCAGAAGTACAACATCTT

Note: the underlined part comes from the synthetic fragment.

##### The pegRNA cassettes in pU3-ALS-S1 and pU3-GAPDH

OsU3p-tGly-Spacer-sgR-rtT-PBS-HDV-TaU3t

AGTAATTCATCCAGGTCACCAAGTTCTAGGATTTTCAGAACTGCAACTTATTTTATCAAGGAATCTTTAAACATACGAACAGATCACTTAAAGTTCTTCTG
AAGCAACTTAAAGTTATCAGGCTTGCATGGATCTTGGAGGAATCAGATGTGCAGTCAGGGACCATAGCACAAGACAGGCGTCTTCTACTGGTGCTACC
AGCAAATGCTGGAAGCCGGGAACACTGGGTACGTTGGAAACCACGTGATGTGAAGAAGTAAGATAAACTGTAGGAGAAAAGCATTTCGTAGTGGGC
CATGAAGCCTTTCAGGACATGTATTGCAGTATGGGCCGGCCCATTACGCAATTGGACGACAACAAAGTCTAGTATTAGTACCACCTCGGCTATCCACATA
GATCAAAGCTGATTTAAAAGAGTTGTGCAGATGATCCGTGGCAACAAAGCACCAGTGGTCTAGTGGTAGAATAGTACCCTGCCACGGTACAGACCCGG
GTTCGATTCCCGGCTGGTGCANNNNNNNNNNNNNNNNNNNNGTTTTAGAGCTAGAAATAGCAAGTTAAAATAAGGCTAGTCCGTTATCAACTTG
AAAAAGTGGCACCGAGTCGGTGCNNNNNNNNNNNNNNNNNNNNNNNNNNNNNNGGCCGGCATGGTCCCAGCCTCCTCGCTGGCGCCGGCTGG
GCAACATGCTTCGGCATGGCGAATGGGACTTTTTTTTTTGTCCTTCTGTTTTTTTAGTCAGTCTCTTTTTTCAGAAGTACAACATCTT

Note: the underlined part comes from the synthetic fragments.

##### The pegRNA cassettes in p35C-ALS-S1 and p35C-GAPDH

35S-CmYLCV-U6-tGly-Spacer-sgR-rtT-PBS-HDV-polyT-HSPt

ATGGAGTCAAAGATTCAAATAGAGGACCTAACAGAACTCGCCGTAAAGACTGGCGAACAGTTCATACAGAGTCTCTTACGACTCAATGACAAGAAGAA
AATCTTCGTCAACATGGTGGAGCACGACACACTTGTCTACTCCAAAAATATCAAAGATACAGTCTCAGAAGACCAAAGGGCAATTGAGACTTTTCAACA
AAGGGTAATATCCGGAAACCTCCTCGGATTCCATTGCCCAGCTATCTGTCACTTTATTGTGAAGATAGTGGAAAAGGAAGGTGGCTCCTACAAATGCCA
TCATTGCGATAAAGGAAAGGCCATCGTTGAAGATGCCTCTGCCGACAGTGGTCCCAAAGATGGACCCCCACCCACGAGGAGCATCGTGGAAAAAGAA
GACGTTCCAACCACGTCTTCAAAGCAAGTGGATTGATGTGATTGGCAGACATACTGTCCCACAAATGAAGATGGAATCTGTAAAAGAAAACGCGTGAA
ATAATGCGTCTGACAAAGGTTAGGTCGGCTGCCTTTAATCAATACCAAAGTGGTCCCTACCACGATGGAAAAACTGTGCAGTCGGTTTGGCTTTTTCTG
ACGAACAAATAAGATTCGTGGCCGACAGGTGGGGGTCCACCATGTGAAGGCATCTTCAGACTCCAATAATGGAGCAATGACGTAAGGGCTTACGAAA
TAAGTAAGGGTAGTTTGGGAAATGTCCACTCACCCGTCAGTCTATAAATACTTAGCCCCTCCCTCATTGTTAAGGGAGCAAAATCTCAGAGAGATAGTCC
TAGAGAGAGAAAGAGAGCAAGTAGCCTAGAAGTAGTCAAGGCGGCGAAGTATTCAGGCACGTGGCCAGGAAGAAGAAAAGCCAAGACGACGAAA
ACAGGTAAGAGCTAAGCATCTAGAAAGTTGAAAACAATCTTCAAAAGTCCCACATCGCTTAGATAAGAAAACGAAGCTGAGTTTATATACAGCTAGAGT
CGAAGTAGTGATTGAACAAAGCACCAGTGGTCTAGTGGTAGAATAGTACCCTGCCACGGTACAGACCCGGGTTCGATTCCCGGCTGGTGCANNNNNN
NNNNNNNNNNNNNNGTTTTAGAGCTAGAAATAGCAAGTTAAAATAAGGCTAGTCCGTTATCAACTTGAAAAAGTGGCACCGAGTCGGTGCNNN
NNNNNNNNNNNNNNNNNNNNNNNNNNNGGCCGGCATGGTCCCAGCCTCCTCGCTGGCGCCGGCTGGGCAACATGCTTCGGCATGGCGAATGGG
ACTTTTTTTTGATATCTCCGGGGCTAATTGAATATGAAGATGAAGATGAAATATTTGGTGTGTCAAATAAAAAGCTGGTGTGCTTAAGTTTGTGTTTTTTT
CTTGGCTTGTTGTGTTATGAATTTGTGGCTTTTTCTAATATTAAATGAATGTAAGATCTCATTATAATGAATAAACAAATGTTTCTATAATCCATTGTGAATG
TTTTGTTGGATCTCTTCTGCAGCATATAACTACTGTATGTGCTATGGTATGGACTATGGAATATGATTAAAGATAAG

Note: the underlined part comes from the synthetic fragment.

##### The pegRNA cassettes in p2xU3-ALS-WS, pG3R23-OsWS, pR1R4-OsWS, and pL4L3-OsWS

OsU3p-tGly-Spacer-sgR-rtT-PBS-HDV-OsU3t-TaU3p-tMet-Spacer-sgR-rtT-PBS-HDV-TaU3t

AGTAATTCATCCAGGTCACCAAGTTCTAGGATTTTCAGAACTGCAACTTATTTTATCAAGGAATCTTTAAACATACGAACAGATCACTTAAAGTTCTTCTG
AAGCAACTTAAAGTTATCAGGCTTGCATGGATCTTGGAGGAATCAGATGTGCAGTCAGGGACCATAGCACAAGACAGGCGTCTTCTACTGGTGCTACC
AGCAAATGCTGGAAGCCGGGAACACTGGGTACGTTGGAAACCACGTGATGTGAAGAAGTAAGATAAACTGTAGGAGAAAAGCATTTCGTAGTGGGC
CATGAAGCCTTTCAGGACATGTATTGCAGTATGGGCCGGCCCATTACGCAATTGGACGACAACAAAGTCTAGTATTAGTACCACCTCGGCTATCCACATA
GATCAAAGCTGATTTAAAAGAGTTGTGCAGATGATCCGTGGCAACAAAGCACCAGTGGTCTAGTGGTAGAATAGTACCCTGCCACGGTACAGACCCGG
GTTCGATTCCCGGCTGGTGCAGGGTATGGTGGTGCAATGGGGTTTTAGAGCTAGAAATAGCAAGTTAAAATAAGGCTAGTCCGTTATCAACTTGAA
AAAGTGGCACCGAGTCGGTGCAAACCTATCTTCTAATTGCACCACCATGGCCGGCATGGTCCCAGCCTCCTCGCTGGCGCCGGCTGGGCAACATGC
TTCGGCATGGCGAATGGGACTTTTTTTTTTCGTTTTGCATTGAGTTTTCTCCGTCGCATGTTTGCAGTTTTATTTTCCGTTTTGCATTGAAATTTCTCCG
TCTCATGTTTGC**AG**CGTGTTCAACATGAATCCAAACCACACGGAGTTCAAATTCCCACAGATTAAGGCTCGTCCGTCGCACAAGGTAATGTGTGAAT
ATTATATCTGTCGTGCAAAATTGCCTGGCCTGCACAATTGCTGTTATAGTTGGCGGCAGGGAGAGTTTTAACATTGACTAGCGTGCTGATAATTTGT
GAGAAATAATAATTGACAAGTAGATACTGACATTTGAGAAGAGCTTCTGAACTGTTATTAGTAACAAAAATGGAAAGCTGATGCACGGAAAAAGG
AAAGAAAAAGCCATACTTTTTTTTAGGTAGGAAAAGAAAAAGCCATACGAGACTGATGTCTCTCAGATGGGCCGGGATCTGTCTATCTAGCAGGCA
GCAGCCCACCAACCTCACGGGCCAGCAATTACGAGTCCTTCTAAAAGCTCCCGCCGAGGGGCGCTGGCGCTGCTGTGCAGCAGCACGTCTAACATT
AGTCCCACCTCGCCAGTTTACAGGGAGCAGAACCAGCTTATAAGCGGAGGCGCGGCACCAAGAAGCGAACAACAAAACAACAAATCAGAGTGGC
GCAGCGGAAGCGTGGTGGGCCCATAACCCACAGGTCCCAGGATCGAAACCTGGCTCTGATACCTTGAATGCGCCCCCACTTGTTTTAGAGCTAGAA
ATAGCAAGTTAAAATAAGGCTAGTCCGTTATCAACTTGAAAAAGTGGCACCGAGTCGGTGCTGCCTATGATACCAATTGGGGGCGCATTCGGCCGG
CATGGTCCCAGCCTCCTCGCTGGCGCCGGCTGGGCAACATGCTTCGGCATGGCGAATGGGACTTTTTTTTTTGTCCTTCTGTTTTTTTAGTCAGTCTCT
TTTTTCAGAAGTACAACATCTT

Note: the underlined part comes from the synthetic fragments.

##### The pegRNA cassettes in p35C-ALS-WS and pL2L1-OsWS

35S-CmYLCV-U6-tGly-Spacer-sgR-rtT-PBS-HDV-tMet-Spacer-sgR-rtT-PBS-HDV-polyT-HSPt

ATGGAGTCAAAGATTCAAATAGAGGACCTAACAGAACTCGCCGTAAAGACTGGCGAACAGTTCATACAGAGTCTCTTACGACTCAATGACAAGAAGAA
AATCTTCGTCAACATGGTGGAGCACGACACACTTGTCTACTCCAAAAATATCAAAGATACAGTCTCAGAAGACCAAAGGGCAATTGAGACTTTTCAACA

AAGGGTAATATCCGGAAACCTCCTCGGATTCCATTGCCCAGCTATCTGTCACTTTATTGTGAAGATAGTGGAAAAGGAAGGTGGCTCCTACAAATGCCA
TCATTGCGATAAAGGAAAGGCCATCGTTGAAGATGCCTCTGCCGACAGTGGTCCCAAAGATGGACCCCCACCCACGAGGAGCATCGTGGAAAAAGAA
GACGTTCCAACCACGTCTTCAAAGCAAGTGGATTGATGTGATTGGCAGACATACTGTCCCACAAATGAAGATGGAATCTGTAAAAGAAAACGCGTGAA
ATAATGCGTCTGACAAAGGTTAGGTCGGCTGCCTTTAATCAATACCAAAGTGGTCCCTACCACGATGGAAAAACTGTGCAGTCGGTTTGGCTTTTTCTG
ACGAACAAATAAGATTCGTGGCCGACAGGTGGGGGTCCACCATGTGAAGGCATCTTCAGACTCCAATAATGGAGCAATGACGTAAGGGCTTACGAAA
TAAGTAAGGGTAGTTTGGGAAATGTCCACTCACCCGTCAGTCTATAAATACTTAGCCCCTCCCTCATTGTTAAGGGAGCAAAATCTCAGAGAGATAGTCC
TAGAGAGAGAAAGAGAGCAAGTAGCCTAGAAGTAGTCAAGGCGGCGAAGTATTCAGGCACGTGGCCAGGAAGAAGAAAAGCCAAGACGACGAAA
ACAGGTAAGAGCTAAGCATCTAGAAAGTTGAAAACAATCTTCAAAAGTCCCACATCGCTTAGATAAGAAAACGAAGCTGAGTTTATATACAGCTAGAGT
CGAAGTAGTGATTGAACAAAGCACCAGTGGTCTAGTGGTAGAATAGTACCCTGCCACGGTACAGACCCGGGTTCGATTCCCGGCTGGTGCAGGGTAT
GGTGGTGCAATGGGGTTTTAGAGCTAGAAATAGCAAGTTAAAATAAGGCTAGTCCGTTATCAACTTGAAAAAGTGGCACCGAGTCGGTGCAAACC
TATCTTCTAATTGCACCACCATGGCCGGCATGGTCCCAGCCTCCTCGCTGGCGCCGGCTGGGCAACATGCTTCGGCATGGCGAATGGGACAACAAC
AAATCAGAGTGGCGCAGCGGAAGCGTGGTGGGCCCATAACCCACAGGTCCCAGGATCGAAACCTGGCTCTGATACCTTGAATGCGCCCCCACTTGT
TTTAGAGCTAGAAATAGCAAGTTAAAATAAGGCTAGTCCGTTATCAACTTGAAAAAGTGGCACCGAGTCGGTGCTGCCTATGATACCAATTGGGGG
CGCATTCGGCCGGCATGGTCCCAGCCTCCTCGCTGGCGCCGGCTGGGCAACATGCTTCGGCATGGCGAATGGGACTTTTTTTTGATATCTCCGGGGCT
AATTGAATATGAAGATGAAGATGAAATATTTGGTGTGTCAAATAAAAAGCTGGTGTGCTTAAGTTTGTGTTTTTTTCTTGGCTTGTTGTGTTATGAATTTG
TGGCTTTTTCTAATATTAAATGAATGTAAGATCTCATTATAATGAATAAACAAATGTTTCTATAATCCATTGTGAATGTTTTGTTGGATCTCTTCTGCAGCATA
TAACTACTGTATGTGCTATGGTATGGACTATGGAATATGATTAAAGATAAG

Note: the underlined part comes from the synthetic fragment.
